# A large accessory genome, high recombination rates, and selection of secondary metabolite genes help maintain global distribution and broad host range of the fungal plant pathogen *Claviceps purpurea*

**DOI:** 10.1101/2020.05.20.106880

**Authors:** Stephen A. Wyka, Stephen J. Mondo, Miao Liu, Vamsi Nalam, Kirk D. Broders

## Abstract

Pangenome analyses are increasingly being utilized to study the evolution of eukaryotic organisms, which is often governed by variable gene content. While pangenomes can provide insight into polymorphic gene content, inferences about the ecological and adaptive potential of such organisms also need to be accompanied by additional supportive genomic analyses. In this study we constructed a pangenome of *Claviceps purpurea* from 24 genomes and examined the positive selection and recombination landscape of an economically important fungal organism for pharmacology and agricultural research. Together, these analyses revealed that *C. purpurea* has a relatively large accessory genome (∼ 38%) that is likely maintained by high recombination rates (ρ = 0.044) and transposon mediated gene duplication. However, due to observations of relatively low transposable element (TE) content (8.8%) and a lack of variability in genome sizes, prolific TE expansion is likely controlled by these high recombination rates, which may additionally be influencing the overall trend of purifying selection across the genome. Despite this trend, we observed a strong positive selection pressure on secondary metabolite genes, particularly within the ergoline biosynthetic cluster where we also revealed that the *lpsA1* and *lpsA2* genes were the result of a recombination event. These results indicate that secondary metabolites are primary factors affecting the diversification of the species into new ecological niches and help maintain its global distribution and broad host range. These results showcase the use of selection and recombination landscapes to identify mechanisms contributing to pangenome structure and primary factors influencing the evolution of an organism.

**Author Summary:** The use of genomic data to better understand the lifestyle of a pathogen and its relationship with its host has expanded our ability to investigate the evolutionary history of these organisms. This in turn has allowed us to decipher and understand the ambiguity surrounding the true nature of the fungal plant pathogen *Claviceps purpurea*. By combining three different types of broad genomic analyses we identified primary factors affecting the evolution and adaptive potential of this pathogen; particularly a large accessory genome, high recombination rates, and positive selection of genes associated with stress tolerance. These factors likely contribute to the pathogen’s global distribution and broad host range. Furthermore, these findings will influence the direction of future research into optimal control methods.

## Introduction

Pangenomes can provide useful insight into a species distribution and lifestyle through examination of gene functional diversity, abundance, and distribution into core and accessory genomes. These variations often provide fitness advantages and promote adaptive evolution of the organism (Araki *et al.* 2006; Hartmann *et al.* 2018; Brynildsrud *et al.* 2019). In prokaryotes the existence of more open pangenomes (large accessory) has been suggested to be the result of adaptive evolution that allows organisms, with large effective population sizes, to migrate into new ecological niches (McInerney *et al.* 2017). Whereas closed pangenomes (larger core) are found to be associated with more obligate and specialized organisms (McInerney *et al.* 2017). Similar results have been identified in fungal species, where a range of saprotrophic to opportunistic yeasts were found to have accessory genomes representing ∼ 9 – 19% of the genes (McCarthy and Fitzpatrick 2019), while *Zymoseptoria tritici*, a global wheat pathogen, has 40% of genes in the accessory genome (Badet *et al.* 2020). This increase in the *Z. tritici* accessory genome reflects the global distribution of this pathogen that must continuously adapt to overcome new host resistances and multiple cycles of annual fungicide applications (Sánchez-Vallet *et al.* 2018; Badet *et al.* 2020). While the identification of pangenome sizes provide valuable knowledge of polymorphic gene content, which can be used to infer the lifestyle of the species (McInerney *et al.* 2017), a combination of pangenomic and alternative genomic analyses provide a deeper understanding of the primary factors that are contributing to pangenome structure and the adaptive trajectory of the organism.

*Claviceps purpurea* is a biotrophic ascomycete plant pathogen that has a specialized ovarian-specific non-systemic lifestyle with its grass hosts (Píchová *et al.* 2018). Despite the specialized infection pattern, *C. purpurea* has a broad host range of ∼ 400 grass species across 8 grass tribes, including economically important cereal crops such as wheat, barley, and rye and has a global distribution (Píchová *et al.* 2018). However, the mechanisms that underlie the evolutionary success of this species is still understudied. Unlike other pathogens of cereal crops, researchers have been unsuccessful in identifying qualitative resistance (R) genes in crop or wild grass varieties (Menzies and Turkington 2015; Menzies *et al.* 2017; Gordon *et al.* 2020). Menzies *et al.* (2017) did note the potential for a complex virulence and host susceptibility relationship of *C. purpurea* on durum and hexaploid wheat varieties, however, virulence was determined if sclerotia weighed > 81 mg; indicating that *C. purpurea* is able to initiate its biotrophic interaction but might be arrested during the final stages of sclerotia development. During infection the fungus does not induce necrosis or hypersensitive response (host mediated cell death) in its host, instead it actively manages to maintain host cell viability to obtain nutrients from living tissue through a complex cross-talk of fungal cytokinin production (Hinsch *et al.* 2015, 2016; Oeser *et al.* 2017; Kind *et al.* 2018a, 2018b). Furthermore, Wyka *et al.* (2020a) revealed evidence of tandem gene duplication occurring in genes often associated with pathogenicity or evasion of host defenses (effectors), which could implicate their role in the success of the species, however, the factors that were influencing these duplication events remain unclear.

*Claviceps purpurea* is also known for its diverse secondary metabolite profile of ergot alkaloids and pigments (Schardl *et al.* 2013; Tudzynski and Neubauer 2014; Neubauer *et al.* 2016; Flieger *et al.* 2019). Fungal secondary metabolites can play important roles in plant-host interactions as virulence factors but can also increase the fitness of the fungus through stress tolerance (Avalos and Carmen Limon, 2015; Píchová *et al.* 2018; Pusztahelyi *et al.* 2019). It was also recently postulated that the evolution of *C. purpurea* was associated with a host jump and subsequent adaptation and diversification to cooler, more open habitats (Píchová *et al.* 2018; Wyka *et al.* 2020a). In addition, likely due to the toxicity of ergot alkaloids, grass grazing mammals showed avoidance in grazing grass infected with *C. purpurea*, suggesting a potential for beneficial effects for the host plant (Wäli *et al.* 2013). This along with other evidence of neutral to positive effects of infection to host plants (Raybould *et al.* 1998; Fisher *et al.* 2007) suggest that *C. purpurea* is a conditional defensive mutualist (Wäli *et al.* 2013).

In this study, we implement a comprehensive population genomic analysis to gain a deeper understating of factors governing the evolution and adaptive potential of *C. purpurea*. Using 24 isolates, from six countries and three continents, we construct the pangenome and subsequently use single-copy core orthologs to identify genes under positive selection. Full genome alignments were further utilized to estimate population recombination rates and predict recombination hotspots. We observed a large accessory genome likely maintained by a large effective population size and high recombination rates, which subsequently influence an overall trend of purifying selection and likely help defend against TE expansion. In addition, we observed that the *lpsA1* and *lpsA2* genes of the well-known ergoline biosynthetic cluster were likely the result of a recombination event.

## Results

### Pangenome analysis

We constructed a pangenome of *Claviceps purpurea* from 24 isolates representing a collection from three continents and six countries (Table 1). Taking advantage of plentiful isolates available from Canada, we sampled more heavily from different provinces and on different host plants. The principal component and phylogenetic analysis revealed substantial genetic variation among the samples. However, the genetic distances were not correlated with geographic distances, such as LM470 (Canada) and Clav04 (USA) grouping closer to isolates from Europe and the isolate from New Zealand (Additional File 1 Fig. S1). In addition, across Canada and USA, isolates from similar regions rarely clustered together and were often intermixed (Additional File 1 Fig. S1B). These results agree with the results from a multi-locus genotyping of extended samples from Canada and Midwestern USA (Liu *et al. unpublished data*). Previous reports (Wyka *et al.* 2020a) showed that *C. purpurea* isolates had similar genome size (30.5 Mb – 32.1 Mb), genomic GC content (51.6% - 51.8%), TE content (8.42% - 10.87%), gene content (8,394 – 8,824), and BUSCO completeness score (95.5% - 98.0%) (Table 1). The pangenome consisted of 205,354 genes which were assigned to 10,540 orthogroups. We observed 6,558 (62.22%) orthogroups shared between all 24 isolates (core genome), of which 6,244 (59.2%) were single-copy gene clusters, while the remaining core orthogroups, 314 (3%), contained paralogs (2 – 8 paralogs per cluster). The accessory genome consisted of 3,982 (37.78%) orthogroups with 2,851 (27.05%) shared by at least two isolates (but not all) and 1,131 (10.73%) were lineage-specific (singletons) found in only one isolate (Fig. 1, Additional File 2 Table S1). Within the accessory genome (including lineage-specific orthogroups) we observed 592 (5.6%) orthogroups containing paralogs, with some isolates containing > 20 genes per cluster (Fig. 1C, Additional File 2 Table S1).

**Fig. 1.**
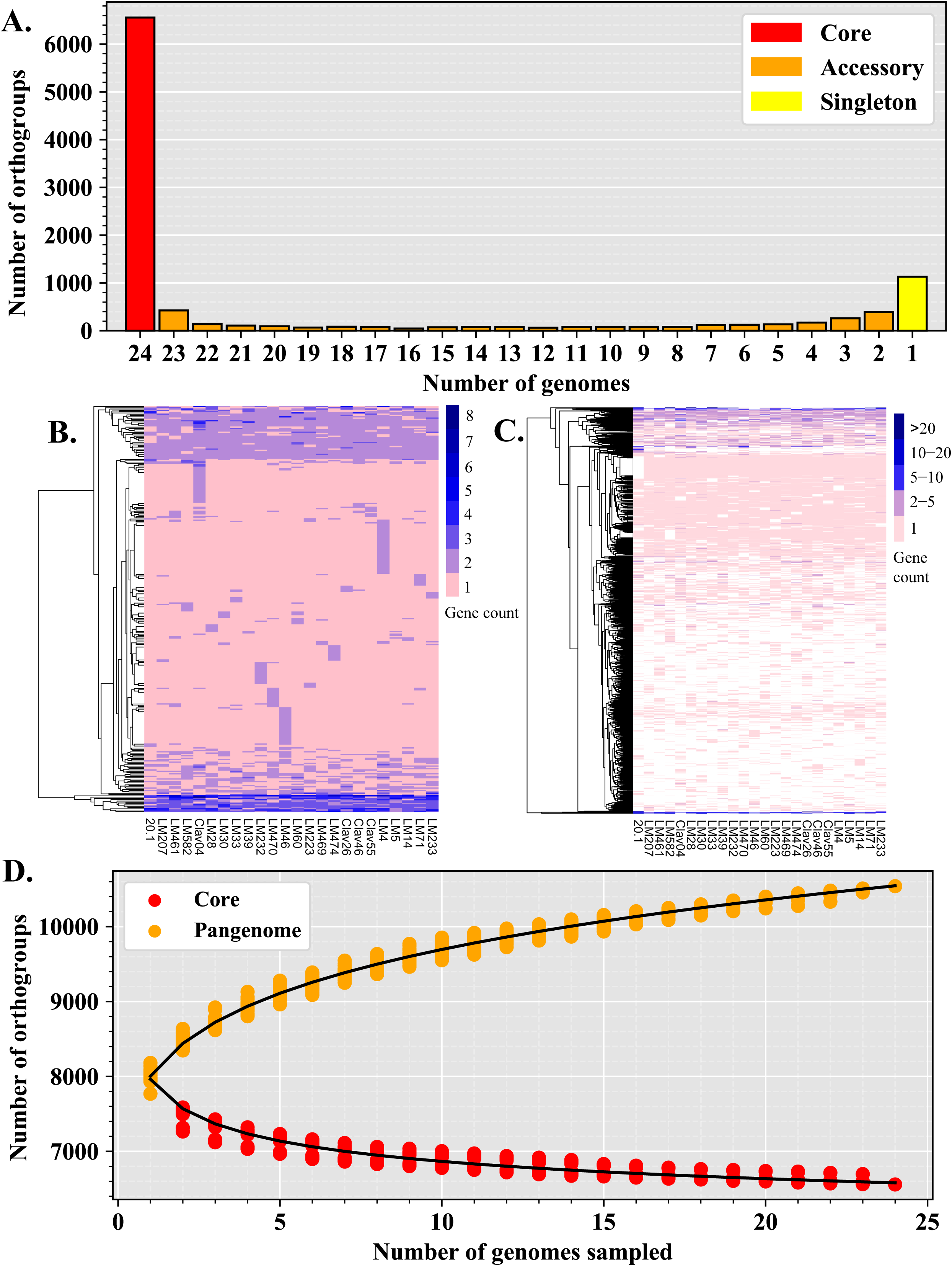
The pangenome of *Claviceps purpurea*. (A) Categorization of orthogroups (gene clusters) into core (shared between all isolates), accessory (shared between ≥ 2 isolates, but not all), and singletons (found in only one isolate) according to the number of orthrogroups shared between genomes. **(B)** Copy number variation in core orthogroups containing paralogs. **(C)** Presence/absence variation and copy number variation of accessory orthogroups, not including singletons. **(D)** Estimation of core and pangenome (core + accessory + singleton) sizes by random resampling of possible combinations of 1 – 24 genomes (dots). Curves were modelled by fitting the power law regression formula: y = Ax^B^ + C.

**Table 1:**
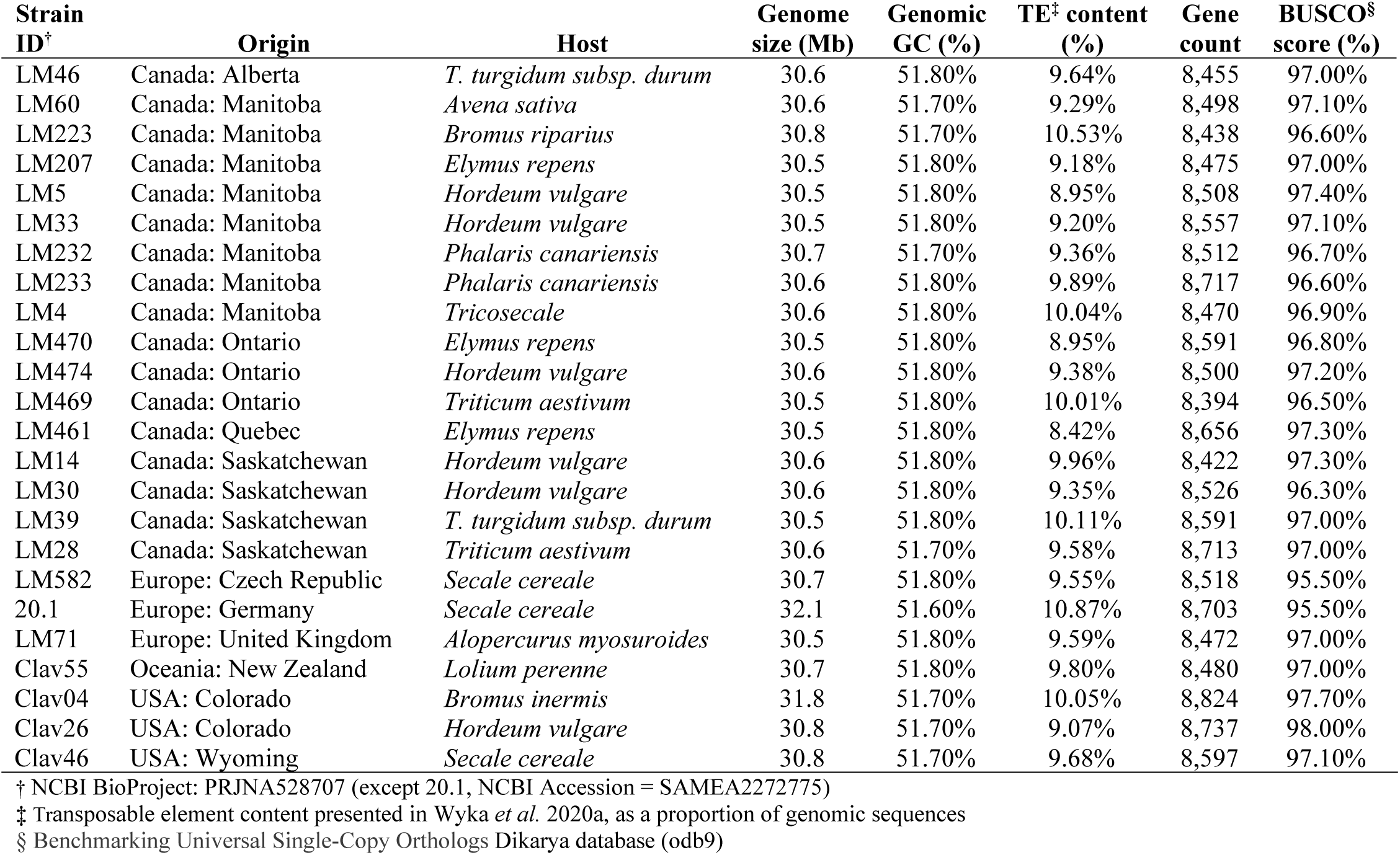
Collection and annotation statistics for the 24 *Claviceps purpurea* genomes used in this study.

| Strain ID<sup>†</sup> | Origin | Host | Genome size (Mb) | Genomic GC (%) | TE<sup>‡</sup> content (%) | Gene count | BUSCO<sup>§</sup> score (%) |
| --- | --- | --- | --- | --- | --- | --- | --- |
| LM46 | Canada: Alberta | *T. turgidum subsp. durum* | 30.6 | 51.80% | 9.64% | 8,455 | 97.00% |
| LM60 | Canada: Manitoba | *Avena sativa* | 30.6 | 51.70% | 9.29% | 8,498 | 97.10% |
| LM223 | Canada: Manitoba | *Bromus riparius* | 30.8 | 51.70% | 10.53% | 8,438 | 96.60% |
| LM207 | Canada: Manitoba | *Elymus repens* | 30.5 | 51.80% | 9.18% | 8,475 | 97.00% |
| LM5 | Canada: Manitoba | *Hordeum vulgare* | 30.5 | 51.80% | 8.95% | 8,508 | 97.40% |
| LM33 | Canada: Manitoba | *Hordeum vulgare* | 30.5 | 51.80% | 9.20% | 8,557 | 97.10% |
| LM232 | Canada: Manitoba | *Phalaris canariensis* | 30.7 | 51.70% | 9.36% | 8,512 | 96.70% |
| LM233 | Canada: Manitoba | *Phalaris canariensis* | 30.6 | 51.80% | 9.89% | 8,717 | 96.60% |
| LM4 | Canada: Manitoba | *Tricosecale* | 30.6 | 51.80% | 10.04% | 8,470 | 96.90% |
| LM470 | Canada: Ontario | *Elymus repens* | 30.5 | 51.80% | 8.95% | 8,591 | 96.80% |
| LM474 | Canada: Ontario | *Hordeum vulgare* | 30.6 | 51.80% | 9.38% | 8,500 | 97.20% |
| LM469 | Canada: Ontario | *Triticum aestivum* | 30.5 | 51.80% | 10.01% | 8,394 | 96.50% |
| LM461 | Canada: Quebec | *Elymus repens* | 30.5 | 51.80% | 8.42% | 8,656 | 97.30% |
| LM14 | Canada: Saskatchewan | *Hordeum vulgare* | 30.6 | 51.80% | 9.96% | 8,422 | 97.30% |
| LM30 | Canada: Saskatchewan | *Hordeum vulgare* | 30.6 | 51.80% | 9.35% | 8,526 | 96.30% |
| LM39 | Canada: Saskatchewan | *T. turgidum subsp. durum* | 30.5 | 51.80% | 10.11% | 8,591 | 97.00% |
| LM28 | Canada: Saskatchewan | *Triticum aestivum* | 30.6 | 51.70% | 9.58% | 8,713 | 97.00% |
| LM582 | Europe: Czech Republic | *Secale cereale* | 30.7 | 51.80% | 9.55% | 8,518 | 95.50% |
| 20.1 | Europe: Germany | *Secale cereale* | 32.1 | 51.60% | 10.87% | 8,703 | 95.50% |
| LM71 | Europe: United Kingdom | *Alopercurus myosuroides* | 30.5 | 51.80% | 9.59% | 8,472 | 97.00% |
| Clav55 | Oceania: New Zealand | *Lolium perenne* | 30.7 | 51.80% | 9.80% | 8,480 | 97.00% |
| Clav04 | USA: Colorado | *Bromus inermis* | 31.8 | 51.70% | 10.05% | 8,824 | 97.70% |
| Clav26 | USA: Colorado | *Hordeum vulgare* | 30.8 | 51.70% | 9.07% | 8,737 | 98.00% |
| Clav46 | USA: Wyoming | *Secale cereale* | 30.8 | 51.70% | 9.68% | 8,597 | 97.10% |

<sup>†</sup> NCBI BioProject: PRJNA528707 (except 20.1, NCBI Accession = SAMEA2272775)
<sup>‡</sup> Transposable element content presented in Wyka *et al.* 2020a, as a proportion of genomic sequences
<sup>§</sup> Benchmarking Universal Single-Copy Orthologs Dikarya database (odb9)

We utilized multiple gene functional categories to get a deeper understanding of how genes of different function were structured within the pangenome. As a proportion of orthogroups within each pangenome category (core, accessory, and singleton) we found that the core genome was significantly enriched in orthogroups that contained genes with conserved protein domains (conserved) (5,471; 84%), transmembrane domains (transmembrane) (1,038; 16%), peptidase and protease domains (MEROPs) (211, 3.2%), and orthogroups of carbohydrate-active enzymes (CAZys) (212, 3.2%) (*P* < 0.01, Fisher’s exact test, Fig. 2A and 2E-G). Effector proteins play major roles in plant-microbe interactions, often conveying infection potential of the pathogen. A total of 257 predicted effector orthogroups were identified; 100 (38.9%) were core, 143 (55.6%) were accessory, and 14 (5.4%) were singletons. Predicted effectors and orthogroups coding for secreted proteins, which also contribute to host-pathogen interactions, were significantly enriched in the accessory genome (143, 5%; 218, 7.6%; respectively) (*P* < 0.01, Fisher’s exact test, Fig. 2C and 2D). Although, the accessory and singleton genomes were largely composed of unclassified orthogroups (1791, 62.8%; 830, 73.4%; respectively) (*P* < 0.01, Fisher’s exact test, Fig. 2H). Lastly, we observed that orthogroups which contained secondary (2°) metabolite genes were similarly represented across all pangenome categories (*P* > 0.05, Fisher’s exact test, Fig. 2B).

**Fig. 2.**
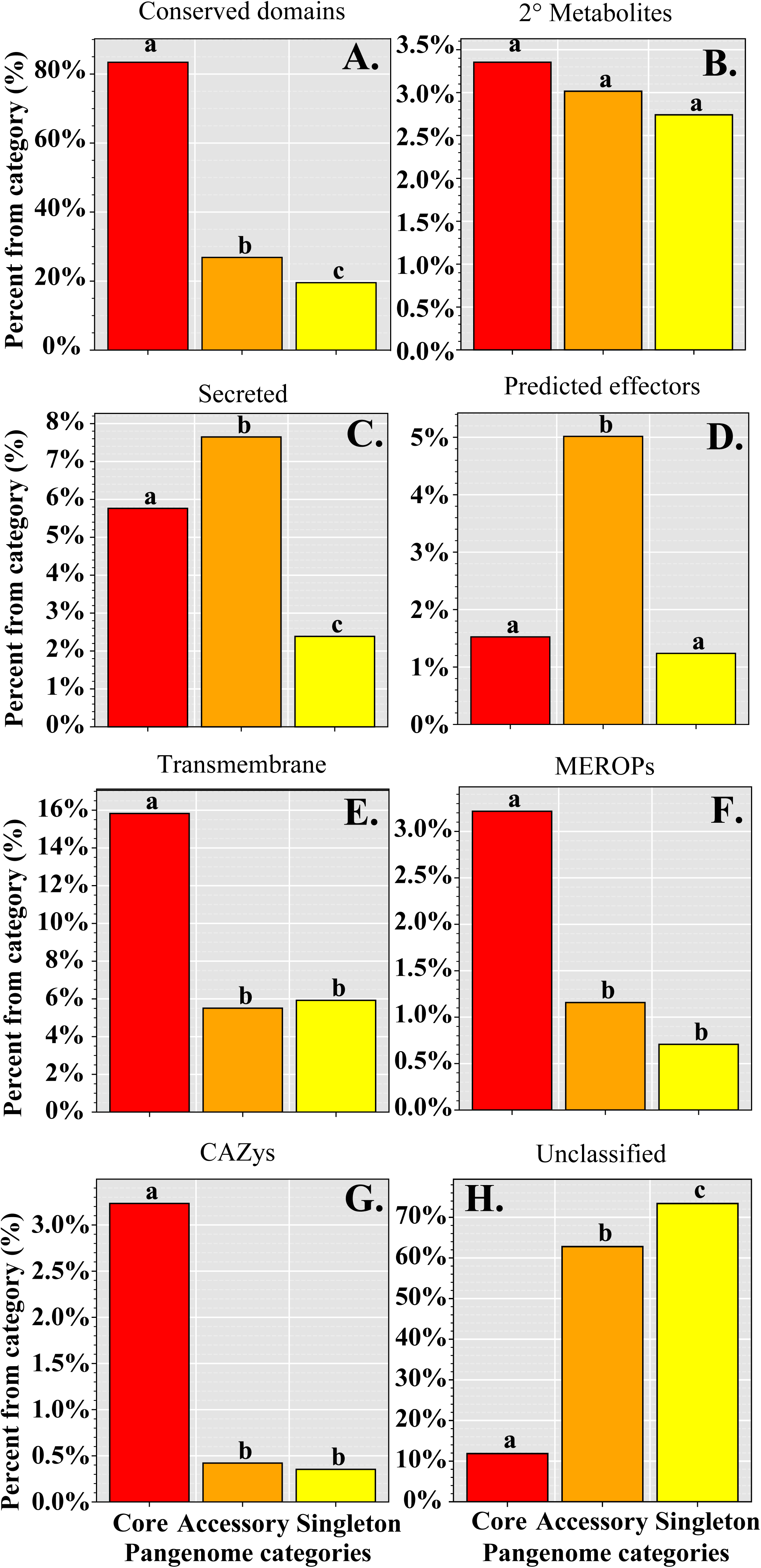
Analysis of predicted protein function across the *Claviceps purpurea* pangenome. Graphs indicate the proportion of orthogroups within each pangenome category of classified protein function. **(A)** Containing conserved protein domains, **(B)** genes found in secondary (2°) metabolite clusters, **(C)** possessing predicted secreted signals, **(D)** predicted to be effectors, **(E)** containing transmembrane domains, **(F)** containing MEROPs domains for proteases and peptidases, **(G)** contain CAZY enzymes, **(H)** all unclassified orthogroups not falling into a previous category. Different letters (within each classification) represent significant differences determined by multi-test corrected Fisher exact test (*P* < 0.01).

As expected, core orthogroups were found to be significantly enriched in general housekeeping and basic cellular functions and development such as protein and ATP binding, nucleus and membrane cellular components, and transmembrane transport, metabolic, and oxidation-reduction processes (Additional File 3 Table S2). Protein domains in core orthogroups were significantly enriched for several WD40-repeat domains, P-loop nucleoside triphosphate hydrolase (IPR027417), armadillo-type fold (IPR016024), and a major facilitator (PF07690) (Additional File 3 Table S2). When narrowing the focus to orthogroups with paralogs, core paralogous orthogroups were enriched in cytochrome P450 domains, and domains associated with trehalose activity (Additional File 3 Table S3). In contrast, the accessory genome was only found to be enriched in a fungal acid metalloendopeptidase domain (MER0001399) and the singleton genome had enrichment for a Tc5 transposase DNA-binding domain (PF03221) (Additional File 3 Table S2). Accessory paralogs were found to be enriched in several protein kinases, Myb-like domains, phosphotransferases, as well as DNA integration and a MULE transposase domain (Additional File 3 Table S3). Overall, our results revealed a large accessory pangenome enriched with genes associated with host-pathogen interactions and an abundance of orthogroups containing paralogs (8.6%), indicating the presence of proliferate gene duplication occurring within the species.

### Positive selection landscape

To further understand the evolution of genes within the pangenome we investigated the positive selection landscape on protein coding genes using 6,244 single-copy core orthologs to compute the ratio of non-synonymous substitutions to synonymous substitutions (dN/dS). Ratios of dN/dS (omega, ω) can provide information of evolutionary forces shaping an organism as genes with ω > 1 may indicate positive or diversifying selection, ω = 1 may indicate neutral evolution, and ω < 1 may indicate negative or purifying selection (Jeffares *et al.* 2015).

Overall, we saw low dN and dS values across all functional categories (Additional File 1 Fig. S3), corresponding to low ω ratios (Fig. 3). This suggests a general trend of purifying selection within *C. purpurea*, although we did identify orthogroups with ω values > 1 (63, 1%), of which 25 (40%) were unclassified (Fig. 3, Additional File 3 Table S4). Notable BLASTp results showed that two conserved genes were related to transcription factors (OG0001193, ω = 1.13, related to subunits Tfc3; OG0004135, ω = 1.21, related to Cys6) and two were related to DNA repair (OG0001034, ω = 1.05, related to mismatch repair PMS1; OG0004027, ω = 1.13, related to XLF (XRCC4-like factor)) (Additional File 3 Table S5). The gene with the highest ω was a transmembrane gene related to a bacteriophage N adsorption protein (OG0001093, ω = 9.79) (Additional File 3 Table S5). Overall, core unclassified genes showed the highest ω values but were not significantly different than predicted effector genes (*P* >> 0.05, multi-test corrected Mann-Whitney U Test, Fig. 3). In contrast, transmembrane, MEROPs, CAZys, and proteins with conserved domains showed the lowest ω values, indicating that these genes are frequently experiencing purifying selection.

**Fig. 3.**
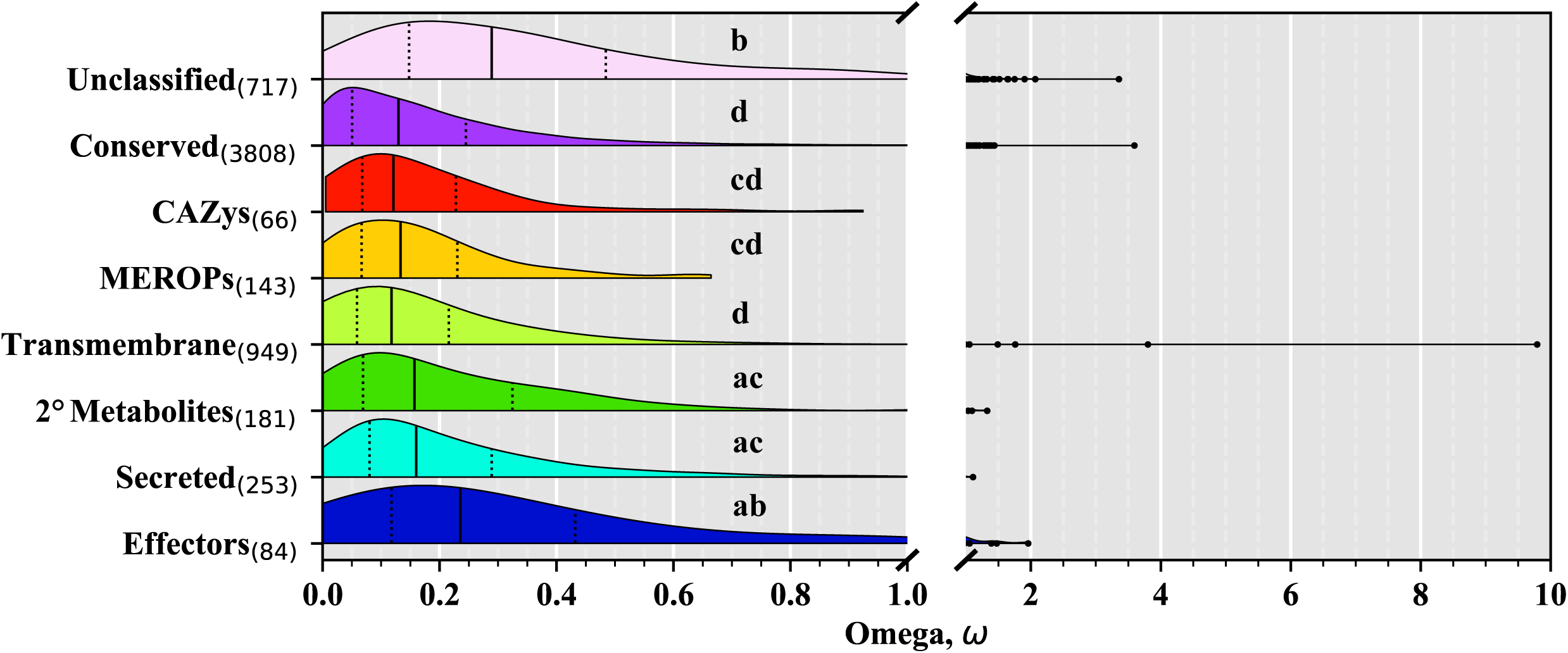
Distribution of omega (ω, dN/dS) ratios within the *Claviceps purpurea* core genome. Violin plots of ω ratios for core single-copy orthogroups protein functional categories. Solid vertical lines within each plot represent the median, while dotted lines represent the 25^th^ and 75^th^ quartile, respectively. Different letters represent significant differences determined by Kruskal-Wallis with *post hoc* multi-test corrected Mann-Whitney U Test (*P* ≤ 0.01).

While ω values, calculated across the entire gene, can provide useful insight on the selective landscape of genes, positive selection and evolution occur at the codon triplet level and can occur in genes where ω, across the entire gene, is < 1 (Goldman and Yang 1994). For this reason, we utilized the CodeML algorithm (Yang 2007a) to more accurately and confidently identify genes with signatures of positive selection. Our results revealed a total of 986 positively selected genes (15.8%) that passed our stringent filtering (Fig. 4A). The majority were genes encoding conserved domains (557, 56.5%) followed by unclassified genes (192, 19.5%). While conserved genes made up the largest portion of genes under putative positive selection, unclassified genes showed the highest proportion of genes with positive selection signatures (26.8%) followed by secondary (2°) metabolite genes (21.5%) (Fig. 4B). We observed an enrichment of positively selected secondary metabolite genes that contained domains for polyketide synthases, several phosphopantetheines, as well as metabolic and catalytic GO terms (*P* ≤ 0.05, Fisher’s exact test, Fig. 4B, Additional File 3 Table S6). In addition, five genes in two known secondary metabolite clusters showed evidence of positive selective signatures; three genes (*easE* ω = 0.51, *lpsB* ω = 0.34, and *lps*C ω = 0.55) in the well-known ergoline biosynthetic cluster (ergot alkaloids) (Schardl *et al.* 2013) and two genes (*tcpC* ω = 0.37 and *tcp*P ω = 0.37) in the epipolythiodiketopiperazine biosynthetic cluster (Dopstadt *et al.* 2016). Within these genes, positive selection was often observed in their AMP-binding and condensation domains but also occurred outside of the domain boundaries (Additional File 3 Table S7). Additionally, one of the three genes responsible for the biosynthesis of fungal cytokinins, a pisatin demethylase cytochrome P450 (Hinsch *et al.* 2015, 2016), had signatures of positive selection (OG0000984, ω = 0.19) (Additional File 3 Table S4). Transmembrane genes saw enrichment of three multicopper oxidase domains (*P* ≤ 0.05, Fisher’s exact test, Additional File 3 Table S6). Of which two transmembrane orthogroups, that contained genes with these domains, also encoded for the laccase CAZy enzymes AA1_1, AA1_2, and AA1_3 (OG0005604, ω = 0.38; OG0002895, ω = 0.22) (Additional File 2 Table S1).

**Fig. 4.**
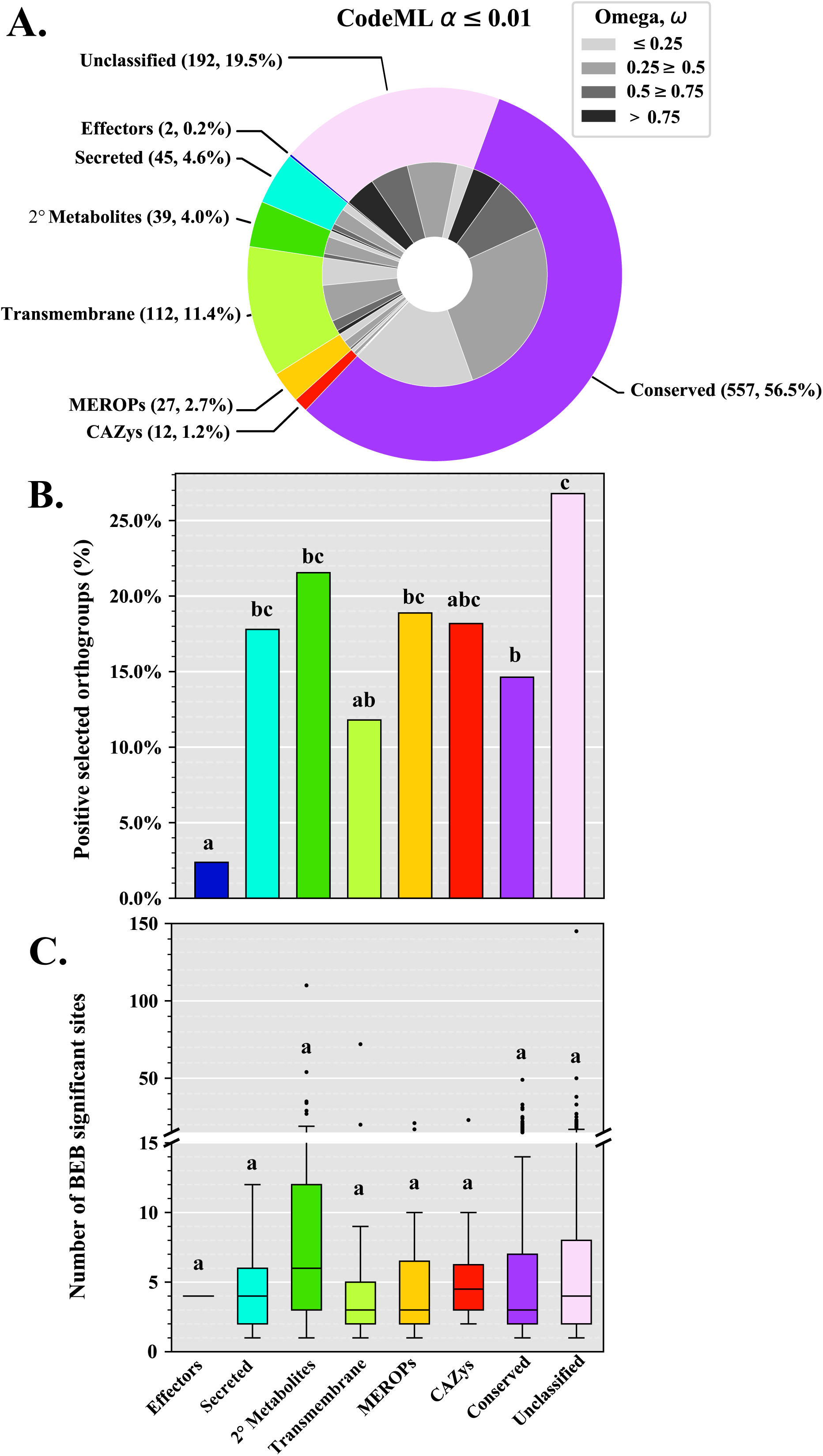
Positive selection landscape within the *Claviceps purpurea* core genome. Positive selection of core single-copy orthogroups protein functional categories as predicted by PAML with the CodeML algorithm. Genes with positive selection signatures were selected after a stringent filtering around an α ≤ 0.01 (*See Methods*). **(A)** The total number of orthogroups in functional categories with signatures of positive selection (outer circle). Omega (ω, dN/dS) ratios of orthogroups within each functional category (inner circle). **(B)** The proportion of orthogroups in each functional category based on the number of orthogroups examined in each category. **(C)** The number of codons with selection signatures in the M8 model of CodeML, as determined by the Bayes Empirical Bayes (BEB) algorithm with an α ≤ 0.01. Different letters **(B, C)** represent significant differences determined by Kruskal-Wallis with *post hoc* multi-test corrected Mann-Whitney U Test (*P* ≤ 0.01). See Additional File 1 Fig. S5 for results from a less stringent filtering of α ≤ 0.05.

There was limited positive selection among predicted core effector genes (Fig. 4B). Only two predicted effector genes (Fig. 4A), corresponding to a proportion of 2.4% of the 84 predicted effector genes examined (Table 2, Fig. 4B), had evidence of positive selection. Suggesting that core effectors might not be under pressure to evolve to overcome host defenses. These two predicted effector genes (OG0003219, ω = 0.76, EffectorP mean score = 0.90 ± 0.028; OG0006565, ω = 1.96, EffectorP mean score = 0.78 ± 0.051) did not have any associated protein domains (Additional File 2 Table S1, Additional File 3 Table S4). We also did not observe any evidence of positive selection in the 10 known virulence factors of *C. purpurea* (Mey *et al.* 2001, 2002; Oeser *et al.* 2002; Scheffer *et al.* 2005a, 2005b; Giesbert *et al.* 2008; Rolke *et al.* 2008; Bormann and Tudzynksi 2009) (Additional File 2 Table S1, Additional File 3 Table S4). In addition, we found no domain enrichment in positively selected secreted genes and CAZys. Peptidase (MEROP) genes only showed enrichment in an alpha/beta hydrolase fold domain (Additional File 3 Table S6).

**Table 2:** PAML and CodeML processing information and filtering of core orthogroups for calculation of dN/dS (ω) ratios and examination of positive selection signatures.

|  |  |  |  |
| --- | --- | --- | --- |
| Total gene clusters (Pangenome) | 10,540 |  |  |
| Single-copy gene clusters | 6,244 |  |  |
| Number of clusters with N/A PAML results | 43 |  |  |
| Cluster Classification (non-redundant) <sup>†</sup>: | Total Pangenome | Total Core<sup>‡</sup> | Single copy<sup>§</sup> |
| Effectors | 257 | 100 (38.9%) | 84 (84.0%) |
| Secreted | 366 | 278 (75.9%) | 253 (91.0%) |
| 2° Metabolites | 313 | 202 (64.5%) | 181 (89.6%) |
| Transmembrane | 1,210 | 998 (82.5%) | 949 (95.1%) |
| MEROPs | 167 | 149 (89.2%) | 143 (96.0%) |
| CAZys | 75 | 68 (90.7%) | 66 (97.1%) |
| Conserved | 4,754 | 3,985 (83.8%) | 3,808 (95.6%) |
| Unclassified | 3,398 | 778 (22.9%) | 717 (92.2%) |

<sup>†</sup> For statistical purposes classification is structured such that each cluster is only represented once (in the order provided), i.e. secreted clusters are those not already classified as effectors, etc.
<sup>‡</sup> Percentage out of total pangenome
<sup>§</sup> Percentage out of total core

Overall, our results revealed a lack of significant positive selection on predicted core effector genes, but a larger proportion of core unclassified and secondary metabolite genes with signatures of positive selection (Fig. 4). It should be noted that secondary metabolite genes also showed the highest number of codons per gene with signatures of positive selection, as determined by the Bayes Empirical Bayes (BEB) algorithm integrated into PAML, however, we did not observe significant differences between gene classifications (Fig. 4C).

### Recombination landscape

Recombination is also an important potential driver of genome evolution and plays a central role in the adaptability of parasitic organisms to overcome host defenses (Morran *et al.* 2011). Our genome-alignments contained 154 of the original 191 scaffolds (Table 3). The 37 missing scaffolds totaled 222,918 bp (average lengths = 6,192 ± 5,676 bp) and corresponded to 59 genes. Thirty-one of the missing scaffolds contained genes that were only part of the accessory genome of which six scaffolds contained two or more genes (Additional File 3 Table S8), suggesting that these scaffolds represent blocks of genetic material that could be lost or gained from isolate to isolate. Most of the genes found on these scaffolds encoded conversed domains associated with either reverse transcriptase, integrases, or helicases (Additional File 3 Table S8), which suggest unplaced repetitive content. Although, one scaffold (scaffold 185) did possess a gene encoding a conserved domain for a centromere binding protein (Additional File 3 Table S8). Together these observations could indicate the potential for dispensable chromosomes, as dispensable and mini-chromosomes often contain higher repetitive content (Peng *et al.* 2019).

**Table 3:**
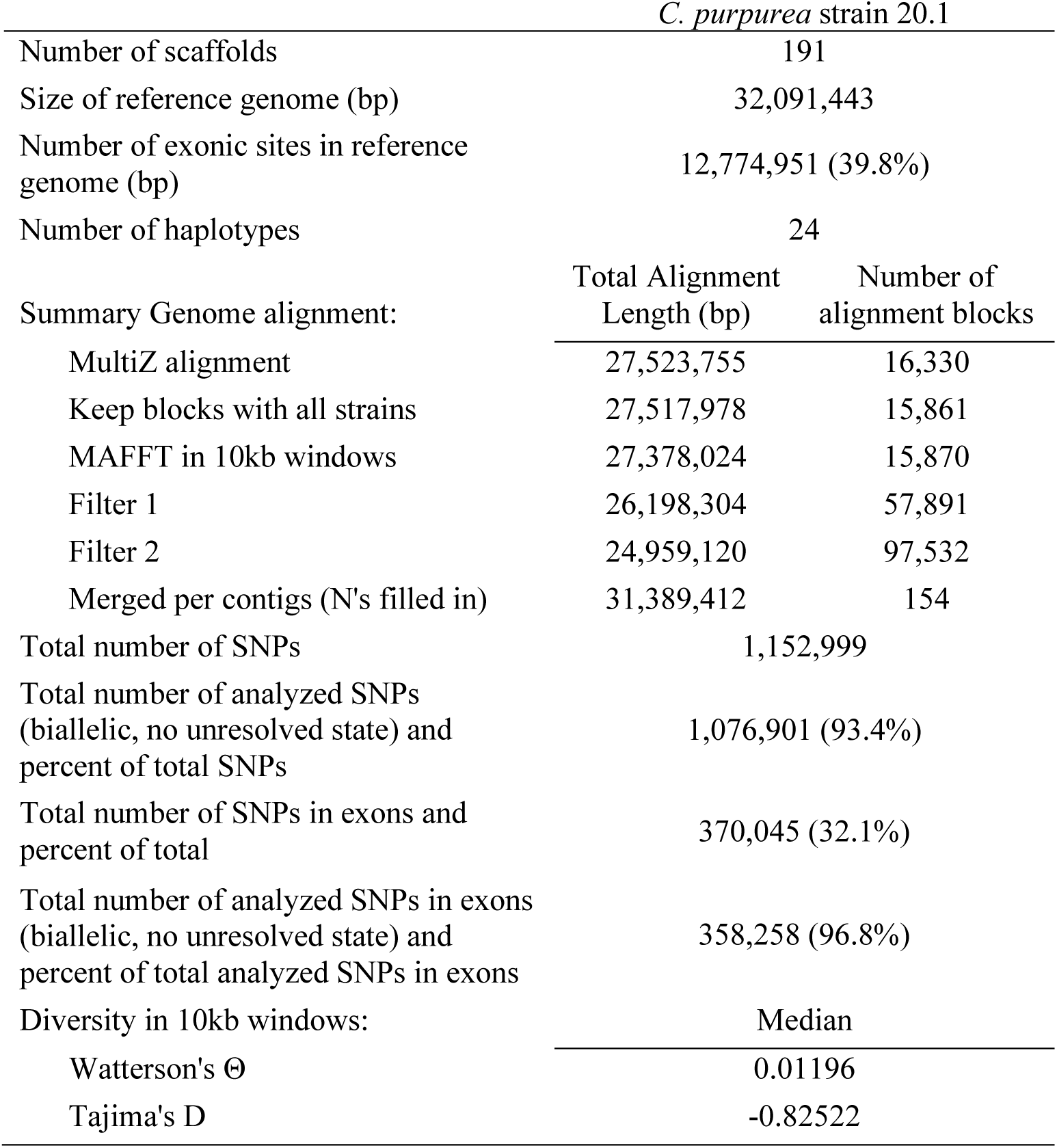
Summary statistics of whole-genome alignment filtering and SNP calls for *Claviceps purpurea*.

From our shared alignments we recovered 1,076,901 biallelic SNPs corresponding to a median nucleotide diversity (Watterson’s θ) of 0.01196 and a Tajima’s D of −0.82522 calculated from 10 kb non-overlapping windows (Table 3). The resulting SNPs were used to infer the population recombination rate (ρ) from the linkage disequilibrium between SNPs based on *a priori* specified population mutation rate θ, which was set to 0.01 based on our nucleotide diversity (Watterson’s θ) (Table 3) (Stukenbrock and Dutheil 2018a). The *C. purpurea* genome recombination landscape was highly variable as some scaffolds showed highly heterogenous landscapes, other scaffolds showed intermixed large peaks of recombination, while others still had more constantly sized peaks across the regions (Fig. 5, Additional File 1 Fig. S6). Overall, the mean genomic population recombination rate in *C. purpurea* was ρ = 0.044. We also examined recombination in specific sequence features and gene type through comparison of mean population recombination rates in exons, introns, 500-bp upstream and downstream of the coding DNA sequence, and intergenic regions based on the annotation of the reference genome (strain 20.1). The distribution of population recombination rates were comparable across different gene features and gene functional categories, although, some significant differences were observed (Fig. 6). In general, we found upstream regions to have the lowest recombination rates, while downstream regions have the highest recombination rates (Fig. 6). The decreased recombination in upstream regions might be the result of mechanisms trying to conserve promotor regions. This trend was observed across different functional gene categories, except in predicted effector genes where exons showed the highest recombination rates and downstream regions with the lowest, although these were not significantly different (Fig. 6B). Across functional categories, secreted genes and transmembrane genes showed the highest recombination rates within each gene feature but were not always significantly different (Fig. 6C).

**Fig. 5.**
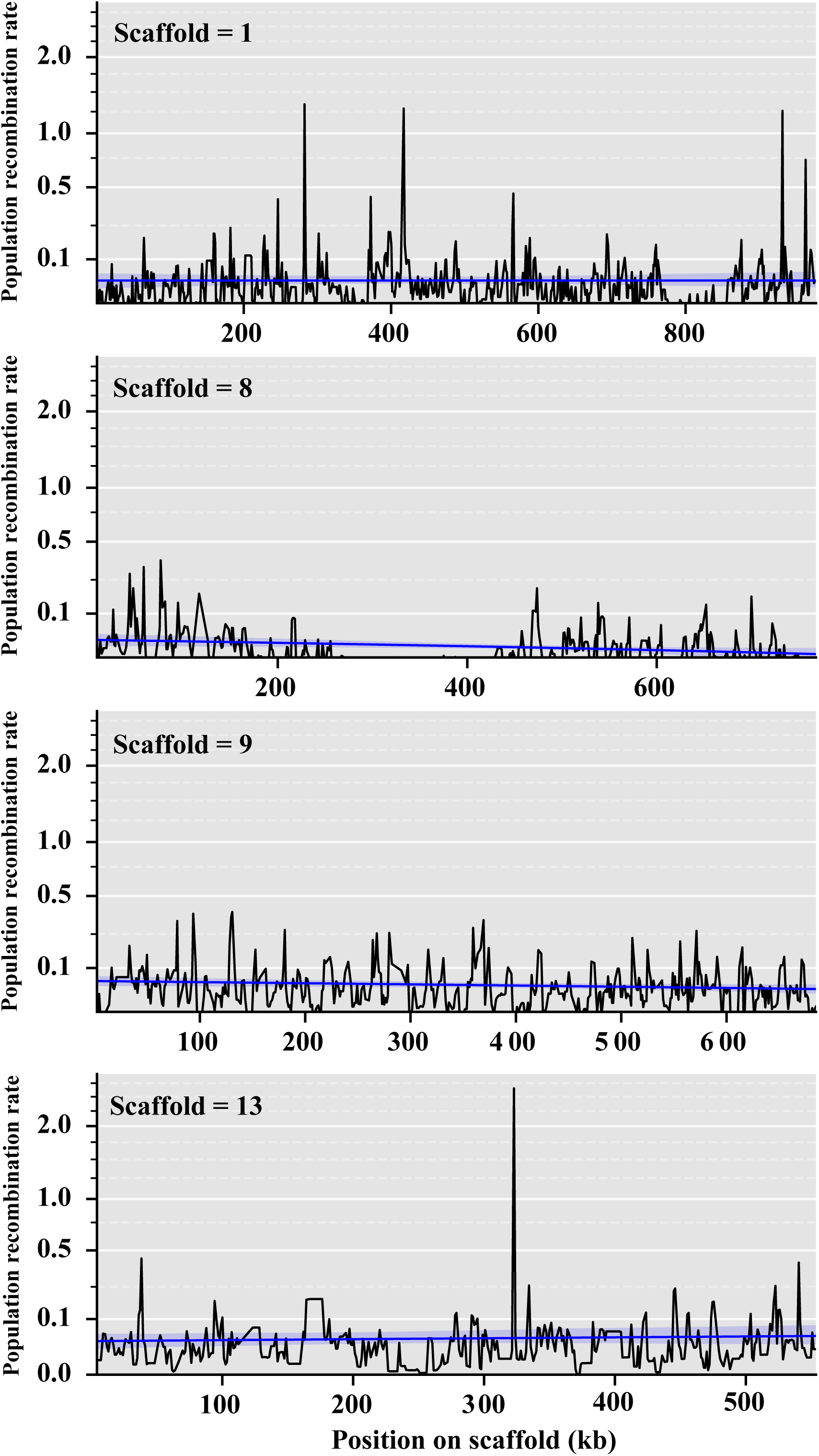
Population recombination rates of representative scaffolds. Estimates of population recombination rates (ρ), in non-overlapping 1 kb windows, across four representative scaffolds displaying the different variation observed across the *Claviceps purpurea* genome. Smoothing curves were calculated from population recombination rates in 10 kb windows. See Additional File 1 Fig. S6 for remaining scaffolds.

**Fig. 6.**
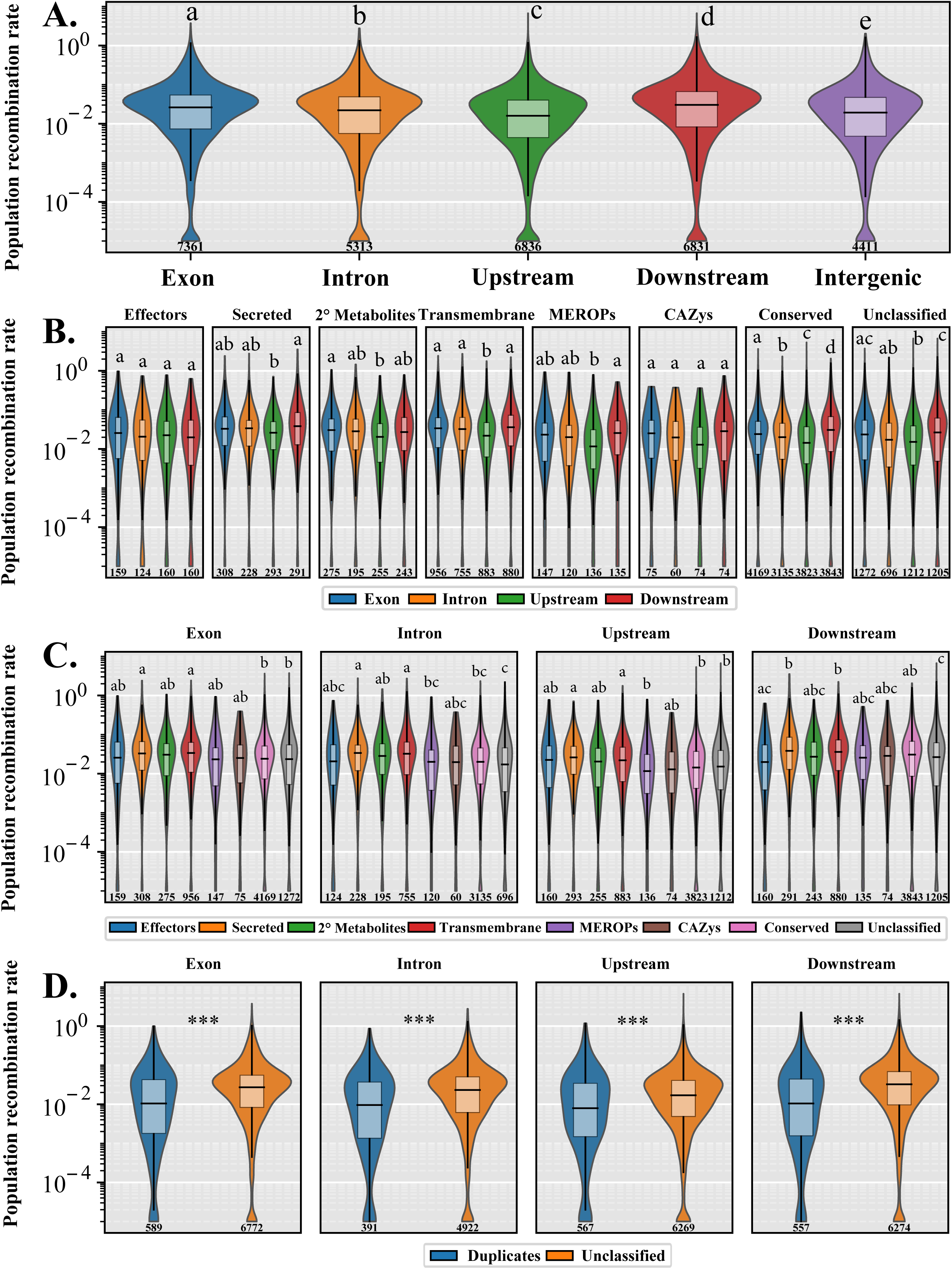
Fine-scale recombination patterns across the *Claviceps purpurea* genome. Plots indicate the distribution of estimated population recombination rates (ρ) between **(A)** different gene features (exons, introns, 500bp upstream and downstream), and **(B-D)** genes of different functional categories and classification. Different letters represent significant differences determined by Kruskal-Wallis with *post hoc* multi-test corrected Mann-Whitney U Test (*P* ≤ 0.01) between data within each plotting window, *** *P* < 0.0001. Sample sizes are embedded below each plot.

Due to the observation of paralogs (Fig. 1) and evidence of tandem gene duplication in *C. purpurea* (Wyka *et al.* 2020a) we investigated the extent recombination might have influenced these events. We found that duplicated genes had lower population recombination rates than all other genes within the genome (Fig. 6D), suggesting that other factors are influencing gene duplication. Due to the absence of RIP (Wyka *et al.* 2020a), transposable elements (TEs) are likely a contributing factor. To investigate the association of duplicated genes with TEs we calculated the average distance of genes to long terminal repeat (LTR) retrotransposons and the average number of flanking LTRs. Results showed duplicated genes were significantly closer to LTRs and had significantly more flanking LTRs than predicted effector and other genes (*P* < 0.0001, multi-test corrected Mann-Whitney U Test, Additional File 1 Fig. S7).

As we observed distinct peaks of recombination (Fig. 5, Additional File 1 Fig. S6), we further utilized LDhot to call statistically significant recombination hotspots by analysis of the intensity of recombination rates in 3 kb (1 kb increments) windows compared to background recombination rates in 20 kb windows (Auton *et al.* 2014; Wall and Stevison 2016; Stukenbrock and Dutheil 2018a). After implementing a cuttoff of ρ ≥ 5 and length of 20 kb (Wall and Stevison 2016) we retained only five recombination hotspots, ranging from 11 kb to 18.5 kb in length (Fig. 7). We observed a recombination hotspot located between the *lpsA1* and *lpsA2* genes of the ergoline biosynthetic cluster, suggesting that this gene duplication event was likely the result of recombination (Fig. 7D). Association of gene functional category and TEs within hotspots varied between region. Some hotspots showed a greater association with duplicated genes and TEs (Fig. 7B-D), while others showed a lower association (Fig. 7A and 7E). In general, genes with conserved protein domains showed the highest presence within hotspots (Additional File 1 Fig. S8). It should be noted that some unclassified genes and genes with conserved protein domains associated with hotspots were also found to be overlapping regions identified as repeats (Fig. 7A-C and 7E). Protein domains found within these genes were associated with ankyrin (IPR002110) and tetratricopeptide (IPR013026) repeats. Only 5 of the 846 duplicated genes (reported in Wyka *et al.* 2020a) found throughout the reference genome were located within predicted recombination hotspots (Fig. 7, Additional File 1 Fig. S8). While Wyka *et al.* (2020a) showed that gene cluster expansion was prevalent among predicted effectors, we only found one non-duplicated predicted effector (CCE30212.1) located within a recombination hotspot (Fig. 7C). Together these results suggest that while recombination may result in important gene duplication, it is not the primary driver of gene duplication within *C. purpurea*.

**Fig. 7.**
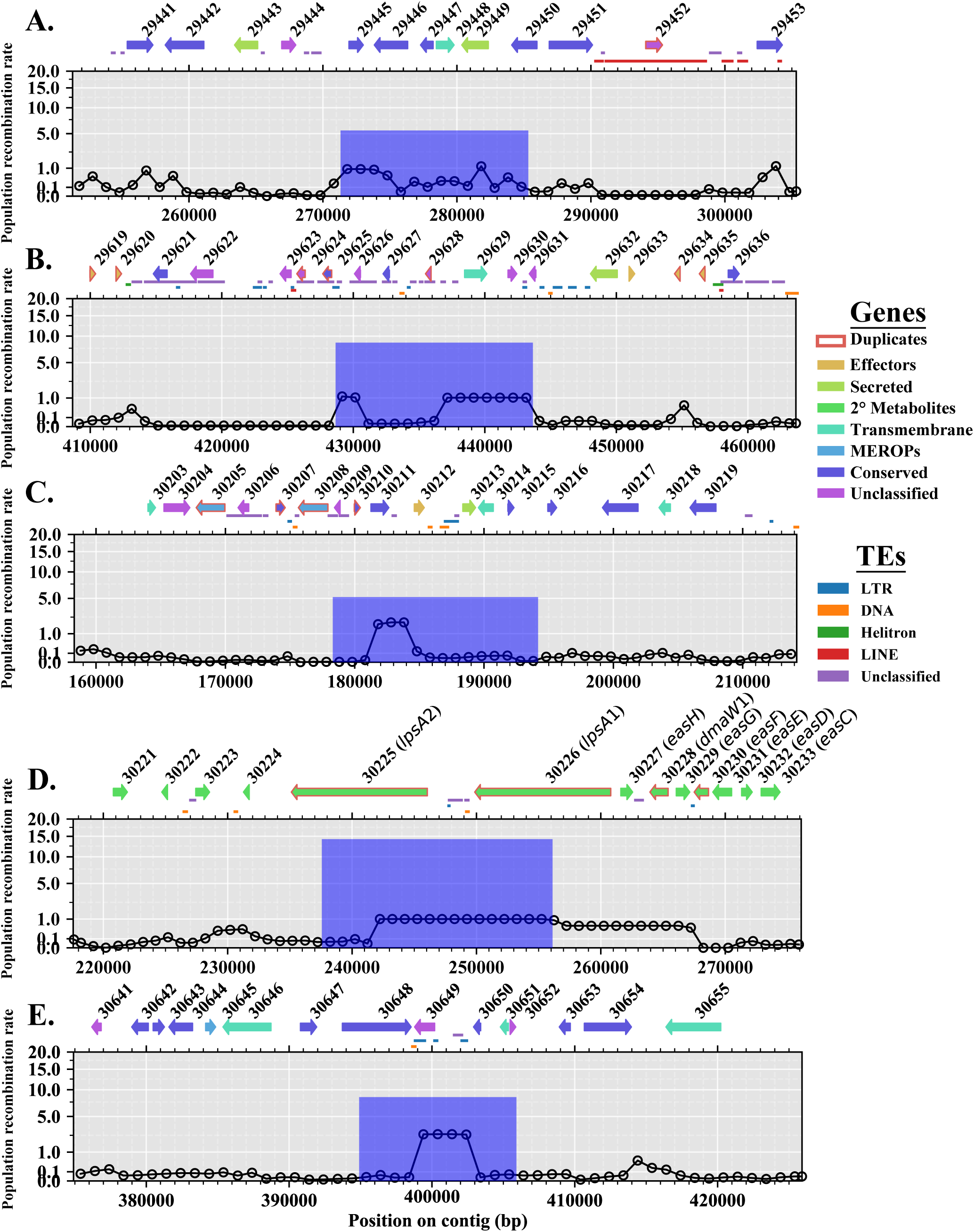
Recombination hotspots predicted in *Claviceps purpurea* with associated genes and transposable elements (TEs). Panels indicate scaffolds: **(A)** scaffold 14; **(B)** scaffold 15; **(C, D)** scaffold 20; **(E)** scaffold 23. Lines indicate background population recombination rates (ρ) estimated in non-overlapping 1 kb windows. Blue bars represent the position, intensity, and width of the predicted hotspots. Genes within the hotspot window and surrounding (± 20 kb) region are depicted by arrows with modified protein ID’s of the reference (strain 20.1; append prefix of “CCE” and suffix of “.1” for protein ID’s) from NCBI. Genes identified as duplicated (≥ 80% identity) from Wyka *et al.* 2020a are outlined in red. TEs are depicted by lines between genes and the corresponding hotspot graph. Colors of arrows and lines correspond to the legend on the right.

## Discussion

Our establishment of a *Claviceps purpurea* pangenome from 24 isolates, as well as, the detection of core genes with signatures of positive selection and analysis of the recombination landscape have provided knowledge into how high recombination rates, gene duplication, and selection of secondary metabolite genes are driving the genomic evolution and adaptation of the species.

The pangenome of *C. purpurea* reveals a large accessory genome with 37.78% accessory orthogroups (27.05% accessory + 10.73% singleton) in comparison to four model fungal pangenomes (*Saccharomyces cerevisiae*, *Candida albicans*, *Cryptococcus neoformans*, and *Aspergillus fumigatus*), which found around 9 – 19% of their genes in the accessory genome (McCarthy and Fitzpatrick 2019). Our results are more comparable to the pangenome of the fungal pathogen *Zymoseptoria tritici* which had an accessory genome comprised of 40% (30% accessory + 10% singleton) of genes (Badet *et al.* 2020). Similar to *C. purpurea*, *Zymoseptoria tritici* is a globally distributed biotrophic fungal pathogen of grasses, notably wheat, suggesting that fungal species with similar life strategies, hosts, and ecological environments could possess comparable pangenome structures as they are under similar evolutionary pressures. Similar factors of lifestyle, effective population size, and habitat have been reported to influence pangenome sizes in bacteria (Mclnerney *et al.* 2017). In fact, *C. purpurea* and *Z. tritici* both experienced enrichment of predicted effector orthogroups in the accessory genome and enrichment of carbohydrate-active enzymes (CAZys) orthogroups in the core genome (Fig. 2) (Badet *et al.* 2020), conveying a comparable similarity between gene functions within pangenome structure regarding the pathogenic lifestyle of these organisms. In addition, Badet *et al.* (2020) suggested that the large accessory genome of *Z. tritici* is likely maintained due to TE activity and a large effective population size as a result of observations of high SNP density, rapid decay in linkage disequilibrium, and high recombination rates (Croll *et al.* 2015; Hartmann *et al.* 2017; Stukenbrock and Dutheil 2018a). The same mechanisms could also explain the large accessory genome observed in *C. purpurea*.

We observed an abundance of orthogroups containing paralogs (8.6%), potentially due to a lack of RIP (Wyka *et al*. 2020a). This presence of gene duplication and association with LTR retrotransposons (Additional File 1 Fig. S7) could be contributing to the large size of the accessory genome, potentially through pseudogenization and/or neofunctionalization. In fact, unclassified genes had the highest ω (dN/dS) ratios (Fig. 3) and the highest proportion of genes with signatures of positive selection (Fig. 4). While this analysis was only conducted on single-copy core genes, it suggests that some of the unclassified accessory genes (Fig. 2H) are undergoing similar evolutionary trends. In addition, the abundance of duplication in accessory unclassified genes (Wyka *et al.* 2020a) and their small sizes (Additional File 1 Fig. S2) can further suggest the presence of pseudogenization and/or neofunctionalization. Badet *et al.* (2020) suggested that TEs were likely contributing to *Z. tritici* accessory genome due to their correlations of TE content with genome size and observations of transcribed TEs. We observed a similar correlation of TE content with genome size (*P* = 0.004, Adj. R^2^ = 0.28), however, our genome sizes and TE content (30.5 Mb – 32.1 Mb, 8.42% - 10.87%, respectively) were not as variable as in *Z. tritici*, which also had a twofold higher TE content (Badet *et al.* 2020). This suggests that TEs play a more important role in *Z. tritici* genome expansion, however, only 0.2% of the orthogroups in *Z. tritici* contained paralogs suggesting that gene duplication is not as common in *Z. tritici* as it is in *C. purpurea* (8.6% paralogs). The lack of gene duplication in *Z. tritici* is likely due to the presence of RIP (Testa *et al.* 2015), which should also reduce TE expansion through silencing (Galagan *et al.* 2003, 2004; Urquhart *et al.* 2018). While we lack RNAseq data to observe TE transcription within *C. purpurea*, observations of TEs with 0% divergence in *C. purpurea* (Wyka *et al.* 2020a) suggest recent TE activity. The observed lack of recombination associated with duplicated genes (Fig. 6D) and association of duplicated genes with LTR transposons (Additional File 1 Fig. S7) would suggest that gene duplication in *C. purpurea* is mediated in part by transposon activity.

Furthermore, we identified 37 missing scaffolds in our population genome alignment with 31 of these containing genes only present in the accessory genome, suggesting the potential for blocks of DNA that could be lost/gained between isolates. Of these accessory scaffolds 15 contained genes encoding conversed domains associated with either reverse transcriptase, integrases, or helicases and one scaffold possessed a gene encoding a conserved domain for a centromere binding protein (Additional File 3 Table S8). Together these could indicate the potential for dispensable mini-chromosomes, as dispensable and mini-chromosomes often contain higher repetitive content (Peng *et al.* 2019). However, even the combination of all 37 missing scaffolds (0.22 Mb) would represent the smallest mini-chromosome known in plant pathogens; 3-fold smaller than *Leptosphaeria maculans* (Balesdent *et al.* 2013), 2-fold smaller than *Nectria haematococca* (Mahmoud and Taga 2012), and 7-fold smaller than *Magnaporthe oryzae* (Peng *et al.* 2019). Many of these scaffolds contained repeated N’s sequences from scaffolding (Schardl *et al.* 2013) and increased repeat content (Additional File 3 Table S8) suggesting that our Illumina based genomes might not have captured the true nature of these scaffolds. Therefore, we did not process these elements further but believe that these are an important aspects of *C. purpurea* evolution and should be a focal point of future research with the advantage of long-read sequencing to more confidently understand their function. Due to these transcriptase rich unplaced scaffolds, the lack of RIP, association of duplicated genes with transposons, and observation of TEs with 0% divergence (Wyka *et al.* 2020a), we believe transposons and/or transcriptases are influencing gene duplication in *C. purpurea*.

Due to the potential for transposon mediated gene duplication, it was remarkable to find relatively low TE content (∼8 - 10%) within *C. purpurea*, especially in the absence of RIP. Other genomic mechanism, such as recombination, may help to limit TE expansion and increases in genome size. Tiley and Burleigh (2015) found a strong negative correlation between global recombination rate, genome size and LTR retrotransposon proportion across 29 plant species, indicating that higher recombination rates actively reduce genome size likely through the removal of LTR elements. A similar function may be affecting LTR content in *C. purpurea*, which would explain the observed differences in LTR content between *Clavicpes* section *Claviceps* (low LTR content, RIP absent) and *Claviceps* sections *Pusillae*, *Paspalorum*, and *Citrinae* (high LTR content, RIP present) (Wyka *et al.* 2020a).

On average we observed a twofold higher mean population recombination rate (ρ = 0.044) in *C. purpurea* than *Z. tritici* (ρ = 0.0217) and tenfold higher than *Z. ardabiliae* (ρ = 0.0045) (Stukenbrock and Dutheil 2018a). As ρ is a function of effective population size and recombination rate per site (ρ = 2*N*_e_ × *r*), these increases could be the result of the increment in recombination rate per site (*r*) and/or effective population size (*N*_e_). Differences in ρ between the two *Zymoseptoria* species was postulated to be due to increased recombination rates per site as it was found that the nucleotide diversity (Watterson’s θ = 2 *N*_e_ x μ, where μ is mutation rate) was 1.6 times higher in *Z. tritici* (0.0139) than *Z. ardabiliae* (0.00866). Under an assumption that both *Z. tritici* and *Z. ardabiliae* have comparable mutation rates, *N*_e_ of *Z. tritici* would only be 1.6 times higher than *Z. ardabiliae*, therefore, the 5 fold higher ρ would likely be caused by higher recombination rates per site (Stukenbrock and Dutheil 2018a). Our observed Watterson’s θ of 0.012 in *C. purpurea* (Table 2) is comparable to *Z. tritici*, suggesting that if mutation rates and effective populations sizes are comparable than the twofold increase in ρ is likely influenced by higher recombination rates per site in *C. purpurea*. Although, *Z. tritici* is a heterothallic organism while *C. purpurea* is homothallic (Esser and Tudzynski 1978) but *C. purpurea* also frequently out-crosses in nature (Amici *et al.* 1967; Tudzynski 2006), suggesting that these factors may provide a difference in effective population sizes between these organisms. In addition, mutation rates might differ between *C. purpurea* and *Z. tritici* for several reasons. Selection pressure associated with agriculture control methods could be driving the mutation of *Z. tritici*, which is subjected to multiple annual fungicide treatments (Torriani *et al.* 2015) and multiple cultivars with various qualitative and quantitative resistance sources (Brown *et al.* 2015). In contrast, control of *C. purpurea* is focused on cultural practices as fungicides have proven inefficient and no resistance crop germplasm has been identified (Menzies and Turkington 2015). While fungicides and crop resistance affect the population structure of *Z. tritici* (Estep *et al.* 2015; Hayes *et al.* 2016; Welch *et al.* 2018), it is plausible to believe they might affect mutation rate or select for strains with a higher mutation or recombination rates. However, we are unaware of any study that has directly examined whether fungicides or crop resistance can have direct or indirect effects on mutation rates. An alternative, and more plausible, hypothesis to explain an increased mutation rate in *Z. tritici* would be associated with the function of RIP, which identifying repeat/duplicated sequences within a genome and introduces C:G to T:A mutations to effectively silence these regions (Galagan *et al.* 2003, 2004; Urquhart *et al.* 2018). It has also been reported that RIP can “leak” into neighboring non-repetitive regions and introduce mutations, thus, accelerating the rate of mutations, particularly those in closer proximity to repeat regions (Fudal *et al.*, 2009; Hane *et al.*, 2015; Van de Wouw *et al.*, 2010). If the mutation rate is increased in *Z. tritici*, either due to RIP “leakage” or selective pressure from fungicides or host resistance the nucleotide diversity in *Z. tritici* could be the result of high mutation rates, whereas the nucleotide diversity in *C. purpurea* could be influenced by higher effective population size and/or recombination rates per site. Higher recombination rates were found to increase the efficacy of purifying selection in both plants (Tiley and Burleigh 2015) and *Z. tritici* (Grandaubert *et al.* 2019). Similarly, *C. purpurea* had an overall trend of purifying selection with skewness towards lower ω values (Fig. 3) and an observed correlation of higher population recombination rates around genes with lower ω ratios (Additional File 1 Fig S9), further suggesting the potential for higher recombination rates in *C. purpurea*.

Additional support, for higher recombination rates per site in *C. purpurea*, could be extrapolated from recombination hotspots, or lack thereof. While we observed evidence of a heterogenous recombination landscapes with several scaffolds showing large peaks in population recombination rates (Fig. 5, Additional File 1 Fig. S6), we only predicted five recombination hotspots (Fig. 7), which is in stark contrast to the ∼1,200 hotspots identified in *Z. tritici* (Stukenbrock and Dutheil 2018b, *Updated dataset*). On average, we did observe higher population recombination rates across scaffolds compared to the rates observed across chromosomes of *Zymoseptoria* (Stukenbrock and Dutheil 2018a), suggesting that the background recombination rate in *C. purpurea* is higher and “flatter”, potentially limiting the detection of hotspots (Auton *et al.* 2014). Overall, this indicates that *C. purpurea* exhibits high recombination rates per site, which potentially helps defend against TE expansion.

While these higher recombination rates are likely influencing the trend of strong purifying selection observed in *C. purpurea*, it might not be the sole factor responsible for the low number of predicted core effector genes with signatures of positive selection (Fig. 4). Wäli *et al.* (2013) classified *C. purpurea* as a conditional defense mutualist with its plant host, as they found that sheep avoided grazing infected grasses and observed that infection rates were higher in grazed pastures compared to ungrazed fields. Other researchers have observed neutral to positive effects of seed set, seed weight, and plant growth on infected plants compared to uninfected plants (Raybould *et al*. 1998; Fisher *et al.* 2007; Wäli *et al.* 2013; Wyka *et al.* 2020 *Unpublished PhD Dissertation*). These factors, along with the broad host range of *C. purpurea* (400+ grass species) and lack of known crop resistance (R) genes, could suggest a lack of strong selection for resistance to *C. purpurea* in grass species (Wäli *et al.* 2013). This could help explain the lack of positive selection observed in predicted core effector genes, implying that effectors are not under strong selection pressure to compete in the evolutionary arms race against host defense. However, it should be noted that positive selection analyses are computed from single-copy core orthologs. Observations of significant enrichment of predicted effector genes in the accessory genome of *C. purpurea* and duplication of effector gene cluster (Wyka *et al.* 2020a) could implicate their role in diversity of infection potential (Sánchez-Vallet *et al.* 2018), however, no host specific races of *C. purpurea* have been identified.

*Claviceps purpurea*, which is suggested to have an ancestral state of plant endophytism (Píchová *et al.* 2018) is also closely related to several mutualistic grass endophytes (i.e. *Epichloe*, *Balansia*, *Atkinsonella*) which have been known to provide beneficial aspects to their hosts mostly through production of secondary metabolites and plant hormones (Clay 1988; Song *et al.* 2016; Xia *et al.* 2018). *Claviceps purpurea* is well-known for its secondary metabolite production and, as we observed, had the second highest proportion of genes with positive selection signatures and the highest number of codons under selection per gene (Fig. 4B and 4C). We also observed two orthogroups with signatures of positive selection containing domains for laccase CAZy enzymes - with some laccases facilitating the biosynthesis of melanin in fungi (Lee *et al.* 2019) - and selection signatures on the cytochrome P450 associated with fungal cytokinin biosynthesis (Hinsch *et al.* 2015). Secondary metabolites are known to increase stress tolerance in fungi (i.e. against UV radiation, oxidative stresses, or colder climates) as has been shown with several groups of pigments, such as melanins and carotenoids (Avalos and Carmen Limon 2015). Therefore, the evolution of secondary metabolites in *C. purpurea* (i.e. ergot alkaloids, ergochromes, or other pigments) can theoretically increase fitness through altering infection potential, stress tolerance, or antimicrobial resistance (Píchová *et al.* 2018; Pusztahelyi *et al.* 2019). The difference in the proportion of secondary metabolites genes under positive selection pressure (such as the ergoline biosynthesis cluster), compared to predicted effectors, indicates that the evolution of secondary metabolite genes in *C. purpurea* is more important to the success of the species than adaptation of core effector proteins. This is in contrast to many fungal plant pathogens of cereal crops, such as *Z. tritici* and the rust fungi in the genus *Puccinia*, that rely on adaptation and diversification of effector proteins for success, particularly due to breeding of crop varieties with R genes (Sánchez-Vallet *et al.* 2018; Badet *et al.* 2020). The selective pressure on secondary metabolites in *C. purpurea* could help explain its evolutionary history as it was recently postulated that evolution of *Claviceps* section *Claviceps*, of which *C. purpurea* resides, occurred tandemly with the radiation of the core Pooideae (Poeae, Triticeae, Bromeae, and Littledaleeae) and was associated with adaptation and diversification to cooler, more open habitats (Kellogg 2001; Sandve and Fjellheim 2010; Píchová *et al.* 2018; Wyka *et al.* 2020a). In addition, the speciation among *C. purpurea* and closely related species demonstrate varied levels of adaptation to ecological niches (Pažoutová *et al.* 2000, 2002, 2015; Douhan *et al.* 2008; Negård *et al.* 2015; Shoukouhi *et al.* 2019; Liu *et al. Submitted*). Similar evolutionary trends towards positive selection of secondary metabolites could be influencing the divergence of these species as well. In fact, all members of *Claviceps* section *Claviceps* had genomes that lack RIP, exhibit gene duplication, and have comparable TE content (Wyka *et al.* 2020a), suggesting that the genomic mechanisms identified in this study might be characteristic of section *Claviceps* as a whole.

## Conclusion

Overall, we observed that the *Claviceps purpurea* pangenome is composed of a large accessory genome that is likely influenced by a large effective population size, high recombination rates, and TE mediated gene duplication. Pseudogenization and neofunctionalization might also be contributing due to the observed TE activity, observations of higher ω ratios, signatures of positive selection in core single-copy unclassified genes, and small size of many accessory unclassified genes. Due to a lack of RIP, prolific TE expansion is likely controlled by high recombination rates, which subsequently may be influencing the overall trend of purifying selection. However, secondary metabolites genes were found to have the highest rates of positive selection on codons within genes, indicating that these genes are a primary factor affecting the diversification of the species into new ecological niches and to potentially help maintain its global distribution and broad host range.

## Materials and Methods

### Genome data

Haploid genome data from a collection of 24 isolates was utilized in this study to provide a comprehensive analysis of *Claviceps purpurea*. The 32.1 Mb reference genomes of *C. purpurea* strain 20.1 was sequenced in 2013 using a combination of single and paired-end pyrosequencing (3 kb fragments) resulting in a final assembly of 191 scaffolds (Schardl *et al.* 2013; NCBI: SAMEA2272775). The remaining 23 isolates were recently sequenced, assembled, and annotated in Wyka *et al.* (2020a: NCBI BioProject: PRJNA528707), representing a collection of isolates from USA, Canada, Europe, and New Zealand (Table 1). The reference genome was subject to an amino acid cutoff of 50 aa to match the other 23 isolates. In this study, we report the pangenome of *C. purpurea*, analysis of the population genomic recombination, and the landscape of genes under positive selection.

Gene functional and transposable element (TE) annotations utilized were those reported in Wyka *et al.* (2020a) and datasets Wyka *et al.* (2020b *Dryad dataset*). In brief, secondary metabolite clusters were predicted using antiSMASH v5 (Blin *et al.* 2019), with all genes belonging to identified clusters classified as “secondary (2°) metabolites”. Functional domain annotations were conducted using InterProScan v5 (Jones *et al.* 2014), HMMer v3.2.1 (Wheeler and Eddy 2013) search against the Pfam-A v32.0 and dbCAN v8.0 CAZYmes databases, and a BLASTp 2.9.0+ search against the MEROPs protease database v12.0 (Rawlings *et al.* 2018). Proteins were classified as secreted proteins if they had signal peptides detected by both Phobius v1.01 (Käll 2007) and SignalP v4.1 (Nielsen 2017) and did not possess a transmembrane domain as predicted by Phobius and TMHMM v2.0 (Krogh *et al.* 2001). Effector proteins were identified by using EffectorP v2.0 (Sperschneider *et al.* 2018) on the set of secreted proteins for each genome. Transmembrane proteins were identified if both Phobius and TMHMM detected transmembrane domains. Transposable elements fragments were identified following procedures for establishment of *de novo* comprehensive repeat libraries set forth in Berriman *et al.* (2018) through a combined use of RepeaModeler v1.0.8 (Smit & Hubley 2015), TransposonPSI (Hass 2010), LTR_finder v1.07 (Xu & Wang 2007), LTR_harvest v1.5.10 (Ellinghaus *et al.* 2008), LTR_digest v1.5.10 (Steinbiss *et al.* 2009), Usearch v11.0.667 (Edgar 2010), and RepeatClassifier v1.0.8 (Smit & Hubley 2015) with the addition of all curated fungal TEs from RepBase (Bao *et al.* 2015). RepeatMasker v4.0.7 (Smit *et al.* 2015) was then used to identify TE regions and soft mask the genomes. These steps were automated through construction of a custom script, TransposableELMT (https://github.com/PlantDr430/TransposableELMT) (Wyka *et al.* 2020a, 2020b).

### Pangenome analysis

The pangenome was constructed using OrthoFinder v2.3.3 (Emms *et al.* 2019), on all genes identified from the 24 genomes, to infer groups of orthologous gene clusters (orthogroups). OrthoFinder was run using BLASTp on default settings. For downstream analysis, gene clusters were classified as secreted, predicted effectors, transmembrane, secondary (2°) metabolites, carbohydrate-degrading enzymes (CAZys), proteases (MEROPs), and conserved domain (conserved) clusters if ≥ 50% of the strains present in a gene cluster had at least one protein classified as such. Gene clusters not grouped into any of the above categories were categorized as unclassified.

Core and pangenome size curves were extrapolated from resampling of 24 random possible combinations for each pangenome size of 1 - 24 genomes and modelled by fitting the power law regression formula: y = Ax^B^ + C using the curve_fit function in the Python module Scipy v1.4.1. These processes were automated through the creation of a custom python script (https://github.com/PlantDr430/FunFinder_Pangenome).

### Positive selection

To investigate the positive selection landscape of genes we collected a total of 6,243 single-copy orthologs across all 24 genomes (See Table 2 for detailed report). For each ortholog cluster sequences were aligned using MUSCLE v3.8.1551 (Edgar 2004) on default settings and values of dN, dS, and dN/dS (omega, ω) were estimated using the YN00 (Yang and Nielsen 2000) method in PAML v4.8 using default parameters. Each ortholog was then individually examined for evidence of positive selection. Guide trees were generated for each ortholog cluster using FastTree version 2.1.10 SSE3 and positive selection was detected using the CodeML algorithm (Yang 2007a) in PAML v4.8 with parameters: NSites = 0 1 2 3 7 8, CodonFreq = 2, seqtype = 1, kappa = 0.3, omega = 1.3, ncatG = 10. Due to high average nucleotide similarities in pairwise BLASTn searches within each ortholog (Additional File 1 Fig. S4) we utilized a stringent filtering method to enhance our confidence in the selection of genes with positive selection signatures. Orthologs were only identified as being under positive selection if they were significant at α ≤ 0.01 using a likelihood ratio test (df −2, χ2 critical value = 9.13) in both the M7 vs. M8 and M2 vs. M1 model comparisons. In addition, orthologs also needed to contain at least one specific amino acid residue significantly (α ≤ 0.01) identified as being under positive selection using the Bayes Empirical Bayes algorithm integrated into PAML (Yang 2007a), in both the M8 and M2 models.

For statistical purposes, each gene cluster was only characterized by one functional category in the order displayed in Table 2 (i.e. secreted genes are those not already classified as effectors, etc). After filtering for positive selection, gene functional categories were examined for enrichment of Pfam, Iprscan, MEROPs, CAZy, and smCOGs domains, as well as, gene ontology (GO) terms (See Methods section *Statistical analyses and plotting*).

### Genome alignment, SNP calling, and recombination

Procedures followed Stukenbrock and Dutheil (2018a), for creation of a fine-scale recombination map of fungal organisms and identification of recombination hotspots. A brief description will be provided below, for a more detailed methodology and explanation of algorithms refer to Stukenbrock and Dutheil (2018a), Auton *et al.* (2014), and Wall and Stevison (2016).

LastZ and MultiZ from the TBA package (Blanchette *et al.* 2004) was used to create the population genome alignment projected against the reference genome, *C. purpurea* strain 20.1 (Schardl *et al.* 2013). Alignments in MAF format were filtered using MafFilter v.1.3.1 (Dutheil *et al.* 2014) following Stukenbrock and Dutheil (2018a). Final alignments were merged according to the reference genome and subsequently divided into nonoverlapping windows of 100 kb. MafFilter was additionally used to compute genome-wide estimates of nucleotide diversity (Watterson’s θ) and Tajima’s D in 10 kb windows. Single nucleotide polymorphisms (SNPs) were called by MafFilter from the final alignment. Principal Component Analysis (PCA) and a Maximum-Likelihood phylogeny were conducted with fully resolved biallelic SNPs (Table 3) using the R package SNPRelate v1.18.1 (Zheng *et al.* 2012) and RAxML v8.2.12 (Stamatakis 2014) using GTRGAMA and 1000 bootstrap replicates, respectively.

The following process was automated through the creation of a custom python script (https://github.com/PlantDr430/CSU_scripts/blob/master/Fungal_recombination.py). LDhat (Auton and McVean 2007) was used to estimate population recombination rates (ρ) from the filtered alignment using only fully resolved biallelic positions. A likelihood table was created for the θ value 0.01, corresponding to the genome-wide Watterson’s θ of *C. purpurea* (Table 3; Julien Dutheil *per comm*), and LDhat was run with 10,000,000 iterations, sampled every 5000 iterations, with a burn-in of 100,000. The parameter ρ relates to the actual recombination rate in haploid organism through the equation ρ = 2*N*_e_ × *r*, where *N*_e_ is the effective population size and *r* is the per site rate of recombination. However, without knowledge of *N*_e_ we cannot confidently infer *r* and thus sought to avoid the bias of incorrect assumptions. Therefore, we reported the population recombination rate (ρ).

Resulting recombination maps were filtered to remove pairs of SNPs for which the confidence interval of the recombination estimate was higher than two times the mean (Stukenbrock and Dutheil 2018a). Average recombination rates were calculated, in regions, by weighing the average recombination estimate between every pair of SNPs by the physical distance between the SNPs. Using the reference annotation file (Schardl *et al.* 2013), we calculated the average recombination rates for features in each gene: 1) exons, 2) introns, 3) 500 bp upstream, and 4) 500 bp downstream with a minimum of three filtered SNPs. Flanking upstream and downstream regions correspond to the 5′ and 3′ regions for forward stranded genes and the 3′ and 5′ regions for reverse stranded genes. We also calculated the average recombination rate for each intergenic region between the upstream and downstream regions of each gene. Introns were added to the GFF3 file using the GenomeTools package (Gremme *et al.* 2013). The original recombination maps produced from LDhat (Julien Dutheil *per comm*) were converted from bp to kb format for use in LDhot (Auton *et al.* 2014) to detect recombination hotspots 1000 simulations and --windlist 10 to create 20 kb background windows (Wall and Stevison 2016). Only hotspots with a value of ρ between 5 and 100 and width < 20 kb were selected for further analysis (Auton *et al.* 2014; Wall and Stevison 2016; Stukenbrock and Dutheil 2018a).

### Statistical and enrichment analyses

Statistics and figures were generated using Python3 modules SciPy v1.3.1, statsmodel v0.11.0, Matplotlib v3.1.1, and seaborn v0.10.0. All multi-test corrections were performed with Benjamini-Hochberg false discovery rate procedure. Enrichment analyses were tested using Fischer’s Exact test with a cutoff α = 0.05. Uncorrected p-values were corrected using Benjamini-Hochberg and Bonferroni multi-test correction with a false discovery rate (FDR) cutoff of α = 0.05. Corresponding p-values from correction tests were averaged together to get a final p-value. Enrichment was performed on protein domain names and GO terms. Orthogroups were only associated with a domain or GO term if ≥ 50% of the strains present in the gene cluster had one gene with the term. This process was automated through creation of a custom python script (https://github.com/PlantDr430/CSU_scripts/blob/master/Domain_enrichment.py).

## Acknowledgements

We would like to thank Julien Dutheil for his assistance in understanding the procedures for estimating fungal recombination rates using LDhat and LDhot.

## Data availability

Most of the relevant data are within the manuscript and supporting files. Additional raw datasets and scripts are available on Dryad: Wyka, Stephen *et al.* (2020), A large accessory genome, high recombination rates, and selection of secondary metabolite genes help maintain global distribution and broad host range of the fungal plant pathogen *Claviceps purpurea*, v1, Dryad, Dataset, doi: 10.5061/dryad.6hdr7sqxp.

## Author Contributions

The project was conceived and designed by S.A.W., S.J.M., and K.B.; S.A.W. performed the research, bioinformatic and software workflows, and analyzed and visualized the data with technical troubleshooting from S.J.M.; M.L., V.N., and K.B. provided management, supervision, research advice, and editorial contributions; S.A.W. wrote the paper with contributions from all other authors.

## Supplemental Figure Captions

Additional File 1 Fig S1. Genetic diversity of 24 *Claviceps purpurea* isolates.

Additional File 1 Fig S2. Average protein lengths (aa) of all orthogroups in *Claviceps purpurea* pangenome.

Additional File 1 Fig S3. Distributions of mean non-synonymous (dN) and synonymous (dS) substitution rates of core single-copy orthogroups in *Claviceps purpurea*.

Additional File 1 Fig S4. Distributions of mean nucleotide identity (%) of core single-copy orthogroups in *Claviceps purpurea*.

Additional File 1 Fig S5. Positive selection landscape within the *Claviceps purpurea* core genome.

Additional File 1 Fig S6. Estimated population recombination rates of *Claviceps purpurea* scaffolds.

Additional File 1 Fig S7. Distributions of genes and their association (distance and flanking counts) to LTR transposable elements.

Additional File 1 Fig S8. Association of genes within recombination hotspots.

Additional File 1 Fig S9. Correlation of recombination rates and omega ratios.

Additional File 2 Table S1. *Claviceps purpurea* pangenome spreadsheet.

Additional File 3 Table S2. Enrichment of protein domains within pangenome.

Additional File 3 Table S3. Enrichment of protein domains within paralogous orthogroups.

Additional File 3 Table S4. PAML and CodeML summarized results.

Additional File 3 Table S5. BLAST results of single-copy core orthologs with an ω (dN/dS) ≥ 1.

Additional File 3 Table S6. Enrichment of protein domains of genes with positive selection.

Additional File 3 Table S7. Positive selection sites within five genes from two known *Claviceps purpurea* biosynthetic clusters.

Additional File 3 Table S8. Annotation information of missing reference scaffolds from 24 isolate whole-genome alignment.

