## Additional_File_1_Fig_S1-S9 for "A large accessory genome, high recombination rates, and selection of secondary metabolite genes help maintain global distribution and broad host range of the fungal plant pathogen *Claviceps purpurea*"

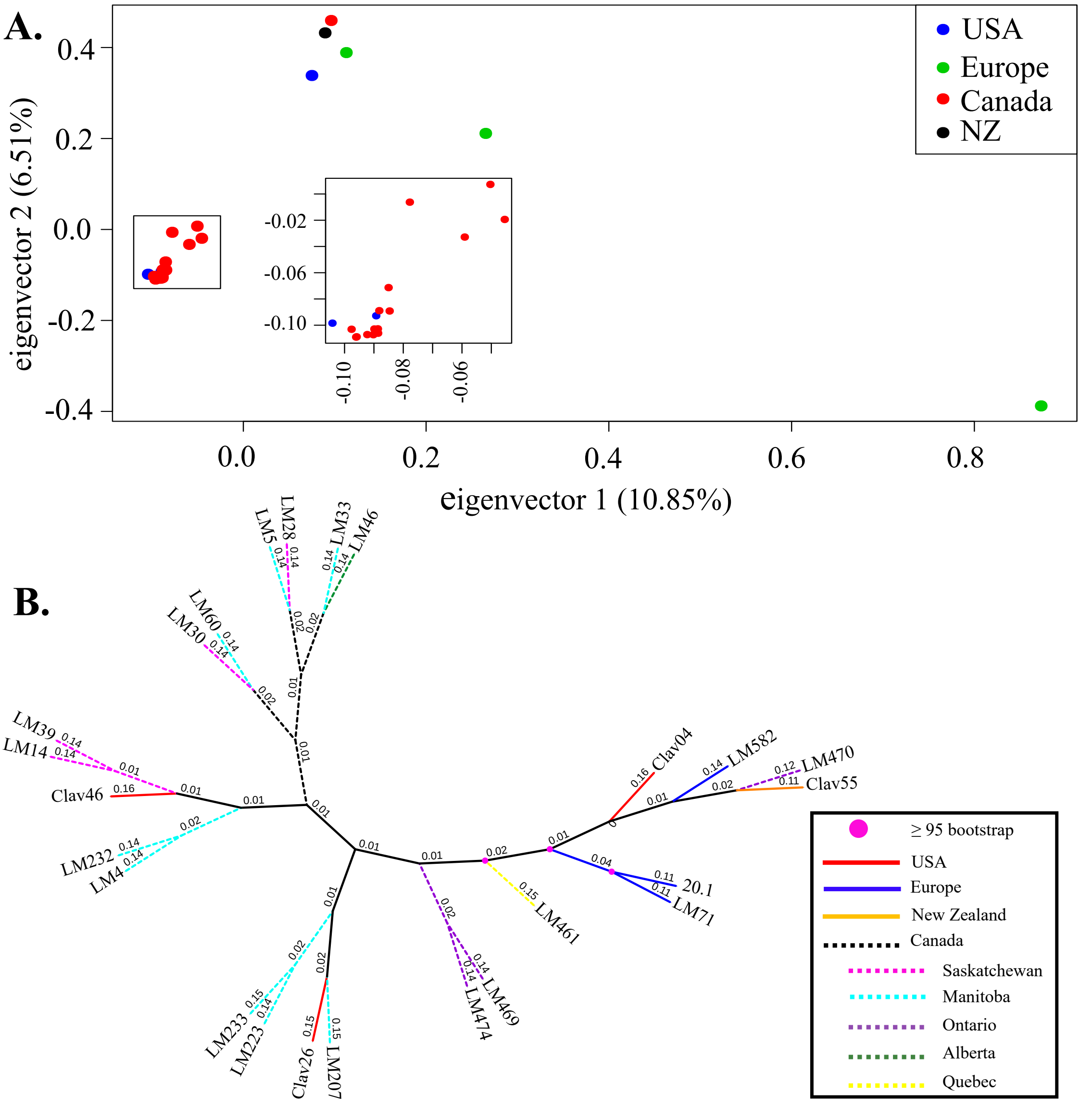


**Fig S1:** Comparison of 24 *Claviceps purpurea* isolates used in this study, based on 1,076,901 bi-allelic SNPs generated from whole genome alignments (Table 2). (**A**) Principal component analysis of SNP matrix with variation explained by each of the axes shown in parentheses. (**B**) Maximum likelihood phylogeny of all isolates used in this study. Colors of branches depict geographical location and pink dots at nodes represent ≥ 95. Branches were reduced for visual purposes; branch lengths were instead written above each line.

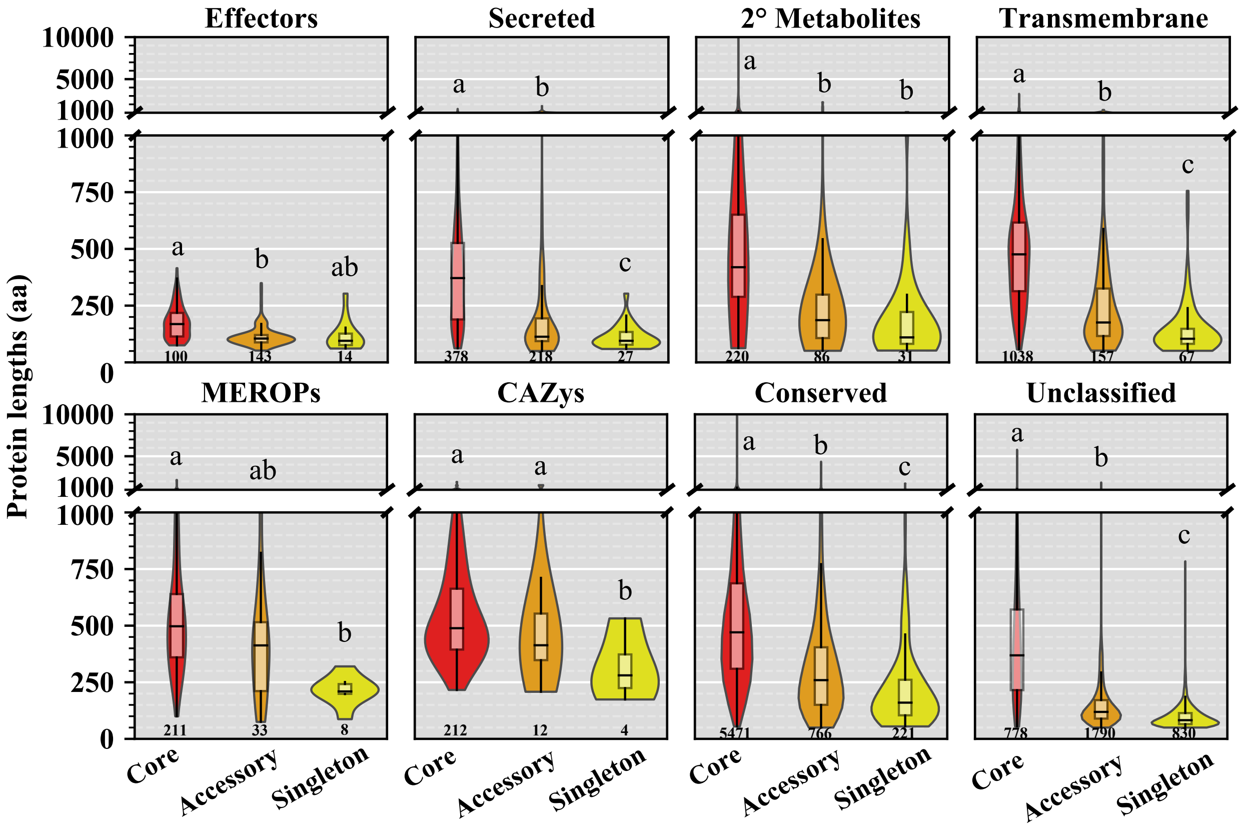


**Fig S2:** Average protein lengths (aa) of all orthogroups identified in each of the *Claviceps purpurea* pangenome categories; core (shared between all isolate), accessory (shared between ≥ 2 isolates, but not all), and singletons (found in only one isolate). Orthogroups are categorized by predicted protein function if ≥ 50% of the isolates present in the orthogroups had one gene identified as such. Different letters represent significant differences determined by Kruskal-Wallis with post hoc multi-test corrected Mann-Whitney U Test ($\alpha \leq 0.01$), with each functional category.

**Fig S3:** Boxplot distributions of mean non-synonymous (dN) and synonymous (dS) substitution rate of core single-copy orthogroups in *Claviceps purpurea*, for each predicted functional category. Orthogroups are categorized by predicted protein function if ≥ 50% of the isolates present in the orthogroups had one gene identified as such. Different letters represent significant differences determined by Kruskal-Wallis with post hoc multi-test corrected Mann-Whitney U Test (α ≤ 0.01), across each substitution category (lower case = dN comparison, upper case = dS comparison).

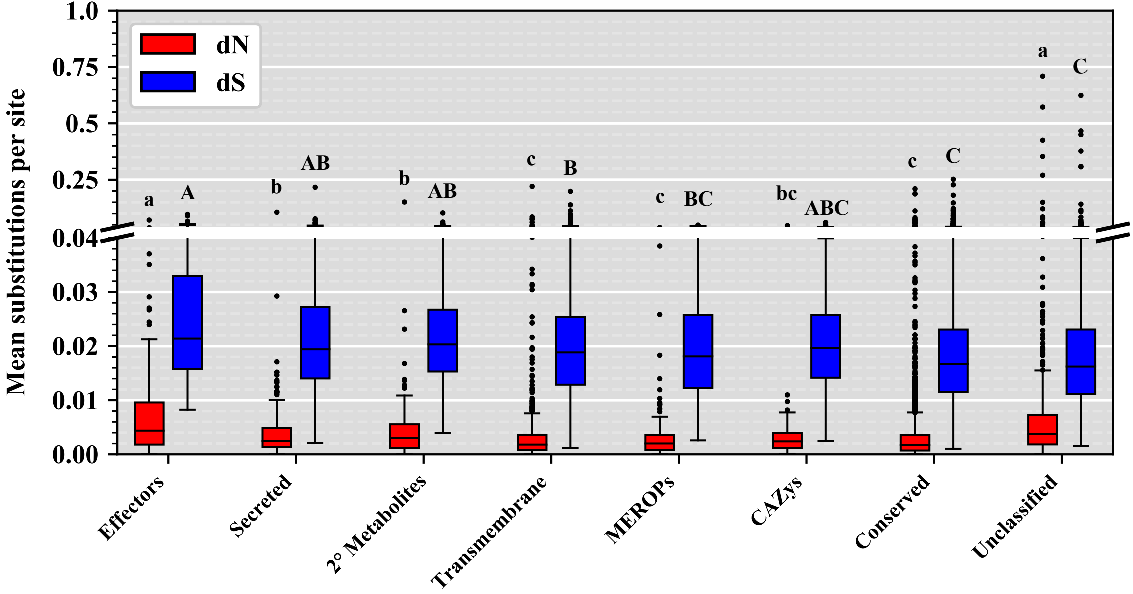

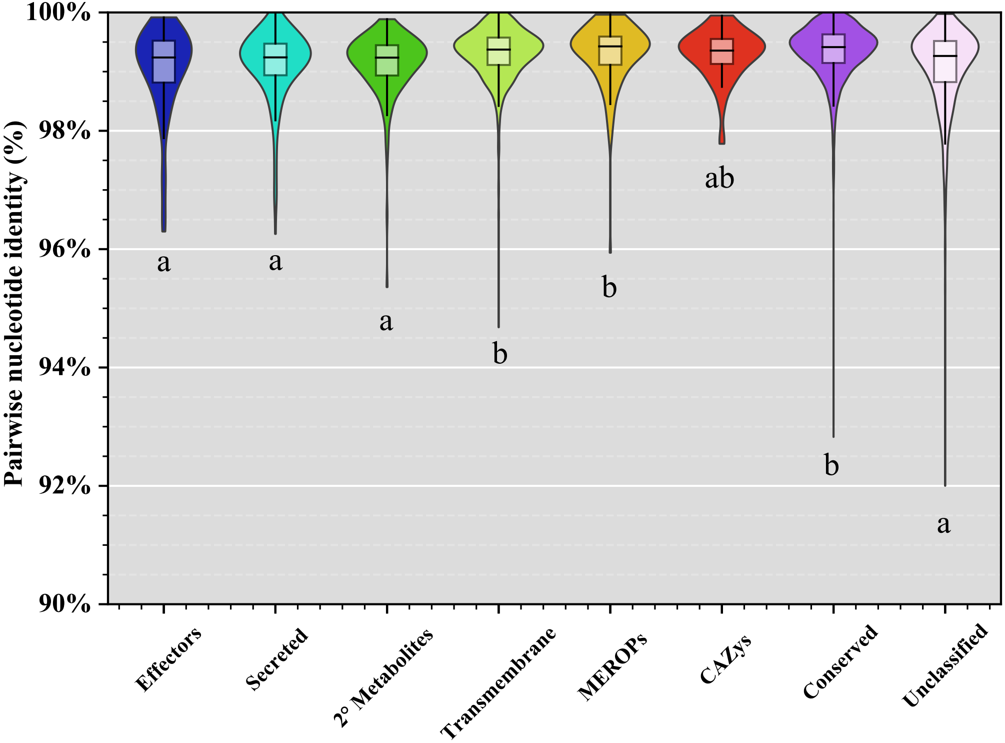


**Fig S4:** Violin plot distributions of mean nucleotide identity (%) of core single-copy orthogroups in *Claviceps purpurea*, for each predicted functional category. Orthogroups are categorized by predicted protein function if ≥ 50% of the isolates present in the orthogroups had one gene identified as such. Different letters represent significant differences determined by Kruskal-Wallis with post hoc multi-test corrected Mann-Whitney U Test (α ≤ 0.01).

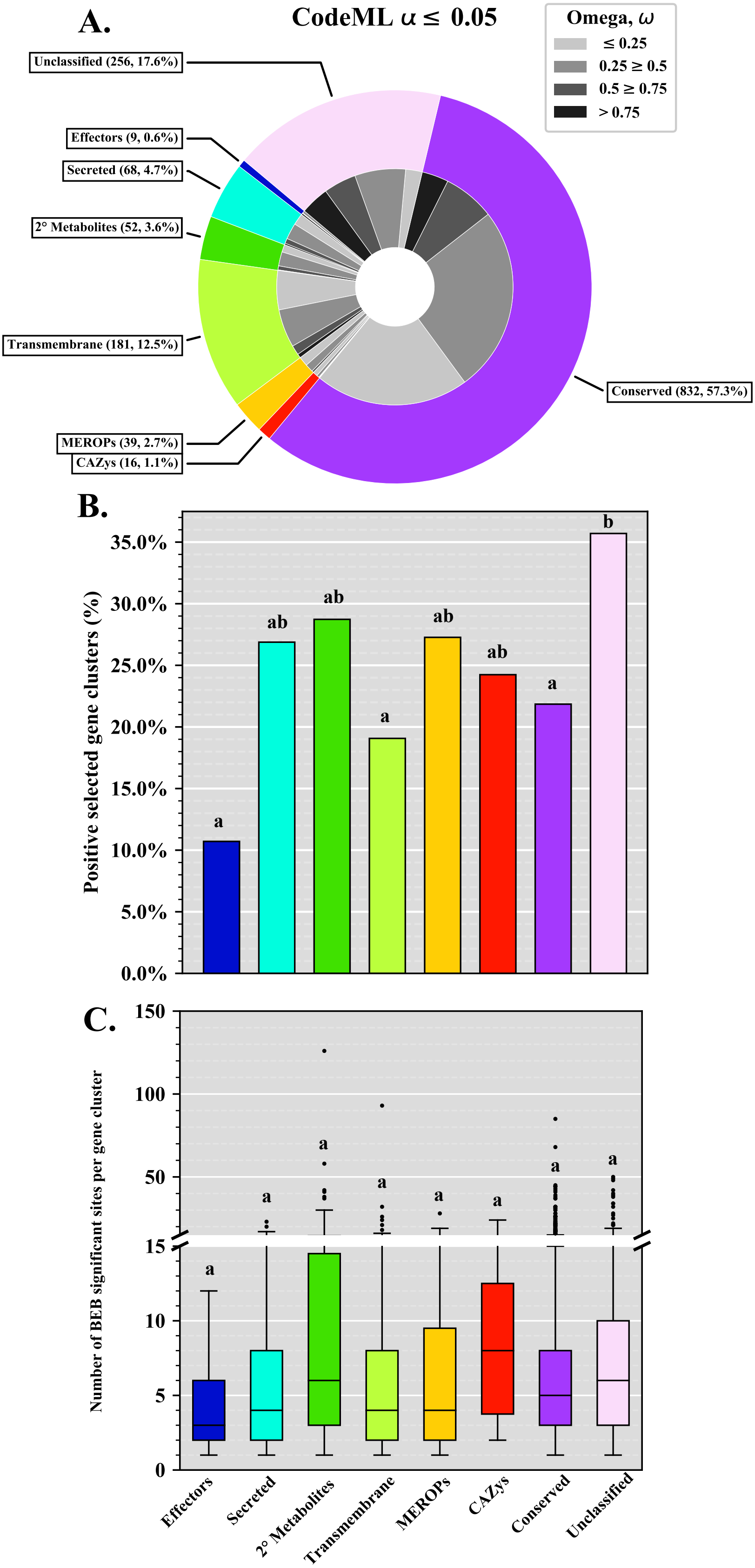


**Figure S5**: Positive selection landscape of core single-copy orthogroups protein functional categories as predicted by PAML with the CodeML algorithm. Genes with positive selection signatures were selected after a stringent filtering around and α ≤ 0.05 (See Methods). A) The total number of orthogroups in functional categories with signatures of positive selection (outer circle). Omega (ω, dN/dS) ratios of orthogroups within each functional category(inner circle). B) The proportion of orthogroups in each functional category based on the number of orthogroups examined in each category. C) The number of codons with selection signatures in the M8 model of CodeML, as determined by the Bayes Empirical Bayes (BEB) algorithm with an α ≤ 0.01. Different letters (B, C) represent significant differences determined by Kruskal-Wallis with post hoc multi-test corrected Mann-Whitney U Test (α ≤ 0.01).

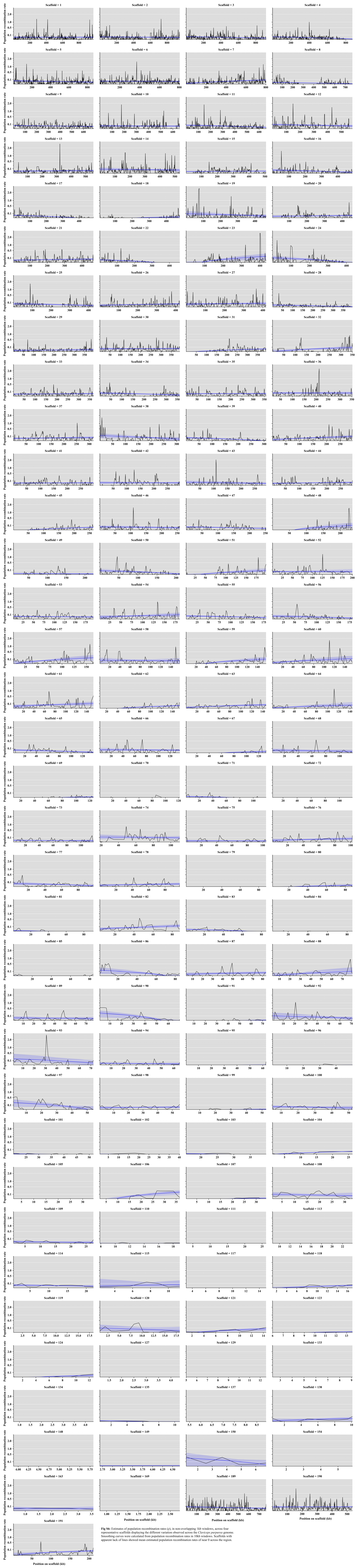


**Fig S6:** Estimates of population recombination rates (ρ), in non-overlapping 1kb windows, across four representative scaffolds displaying the different variation observed across the *Claviceps purpurea* genome. Smoothing curves were calculated from population recombination rates in 10kb windows. Scaffolds with apparent lack of lines showed mean estimated population recombination rates of near 0 across the region.

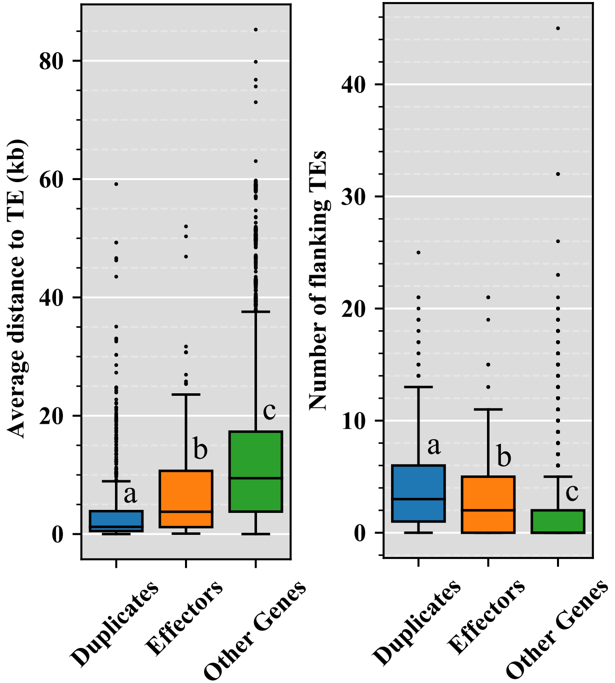


**Fig S7:** Boxplot distributions of putative duplicated genes (≥ 80% identity), predicted effectors, and all other genes in *Claviceps purpurea* showing the mean distance (kb) of each gene to the closest transposable element (TE) fragment (5’ and 3’ flanking distances were averaged together) and the mean number of flanking TE fragments. Different letters represent significant differences determined by Kruskal-Wallis with p*ost hoc* multi-test corrected Mann-Whitney U Test (α ≤ 0.01).

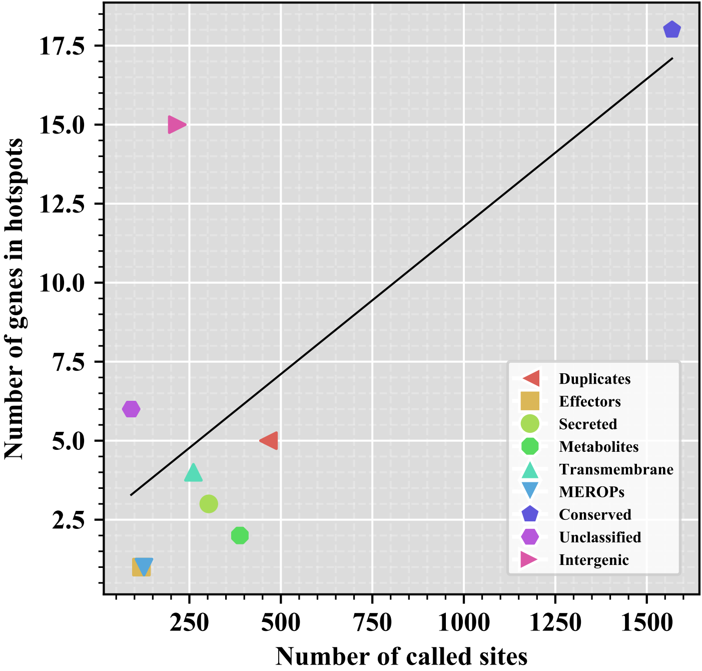


**Fig S8:** Number of genes and intergenic regions within the boundaries of the five predicted recombination hotspots in *Claviceps purpurea* as a function of the number of called sites. Line corresponds to ordinary least square regression.

**Fig S9:** Correlation of estimated population recombination rate to omega, (ω, dN/dS) ratios of all core single-copy orthologs. Points represent median values and error bars indicate the first and third quartiles of each distribution. The x-axis was binned two different ways for clarity of visualization. **A**) Bins with equal point densities. Medians of bins were fit to a power law regression $y = Ax^{-B} + C$. **B**) Bins with unequal densities centered around population recombination rate values of 0, 0.05, 0.1, 0.15, 0.2, 0.3, 0.4, 0.5, 0.6, 0.7, 0.8, 0.9, 1, 1.25, 1.5, 2. Original (non-binned) data was fit to a linear regression, shaded regions depicts 95% confidence interval.

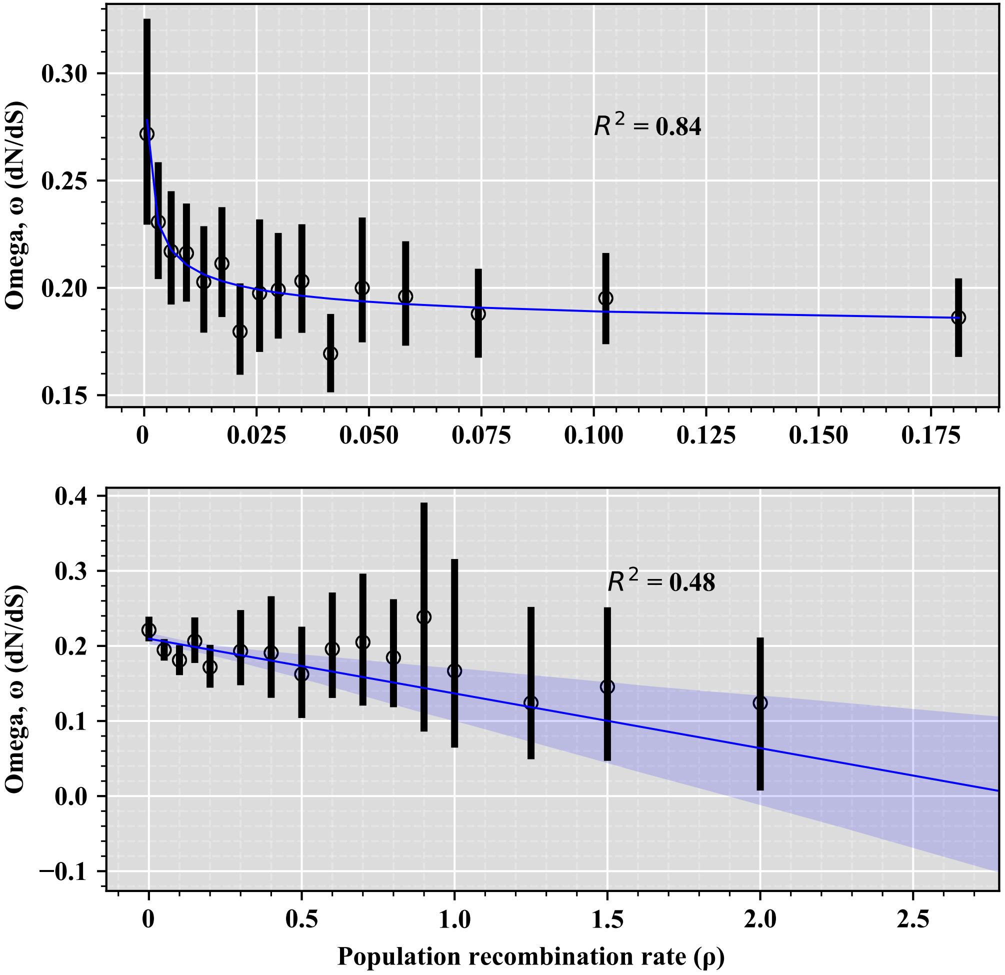
